# Transduction mechanisms for cold temperature in mouse trigeminal and vagal ganglion neurons innervating different peripheral organs

**DOI:** 10.1101/2021.08.05.455314

**Authors:** Katharina Gers-Barlag, Pablo Hernández-Ortego, Eva Quintero, Félix Viana

## Abstract

Thermal signals are critical elements in the operation of interoceptive and exteroceptive neural circuits, essential for triggering thermally-driven reflexes and conscious behaviors. A fraction of cutaneous and visceral sensory endings are activated by cold temperatures. Compared to somatic (DRG and TG) neurons, little is known about the mechanisms underlying cold sensitivity of visceral vagal neurons. We used molecular, pharmacological and genetic tools for a side-by-side characterization of cold-sensitive (CS) neurons in adult mouse trigeminal (TG) and vagal ganglia (VG).

We found that CS neurons are larger in size and more abundant in VG than in TG. In VG, the majority of CS neurons co-express TRPA1 markers and cold-evoked responses are severely blunted in *Trpa1* KO mice. Cold sensitivity was evident in neurons with the highest TRPA1 expression. In contrast, TRPM8 deletion or pharmacological TRPM8 blockade had little impact on VG cold sensitivity. Consistent with these findings, in *Trpm8^eYFP^* reporter mice we found limited expression of TRPM8 in VG and restricted to the rostral jugular ganglion. *In vivo* retrograde labelling of airway-innervating vagal neurons demonstrated their enhanced cold sensitivity and a higher expression of TRPA1 compared to neurons innervating the stomach wall.

In contrast, the majority of CS TG neurons co-express TRPM8 markers and their cold sensitivity is reduced after TRPM8 deletion or blockade. However, pharmacological or genetic reduction of TRPA1 showed that these channels contribute significantly to high-threshold cold sensitivity in TG, suggestive of a role in noxious cold sensing. In both ganglia, a fraction of CS neurons responded to cooling by a mechanism independent of TRPA1 or TRPM8 yet to be characterized.

Finally, in both ganglia, sensitivity to cold varied widely and was enhanced by the potassium channel blocker 4-AP. This effect was independent of the cold sensor expressed by the neuron, suggestive of a common excitability brake mechanism.

**Significance statement:** Temperature sensing and its regulation is a critical homeostatic function. Little is known about the molecular mechanism of cold sensing by visceral sensory endings and their relative weight in different visceral organs. This study highlights important differences in thermotransduction mechanisms between somatic (trigeminal) and visceral (vagal) primary sensory neurons, establishing a critical role of TRPA1 channels in visceral cold transduction. The study describes quantitative differences in cold sensitivity of visceral neurons innervating the stomach and the lower airways, suggesting that cold transduction mechanisms may be fine-tuned to the specific needs of different organs. This study significantly advances our understanding of cold sensing in trigeminal and vagal neurons and reveals distinct drug targets for the pharmacological modulation of these thermoreceptors.

## Introduction

The ability to detect changes in ambient temperature is a critical physiological process. Sensing temperature allows the recognition of pleasurable or potentially harmful thermal stimuli and is also required for the regulation of body temperature, triggering a variety of homeostatic responses and guiding protective behaviors (Morrison & Nakamura, 2019; Senaris et al., 2018).

Cooling the skin surface evokes multiple sensory percepts. As temperature drops, around 15 °C, the sensation of cold turns into aching or burning pain (Viana & Voets, 2020). Single fiber recordings from cutaneous nerves in different species, including humans, identified diverse sensory endings activated by cold temperatures with variable temperature thresholds (Schepers & Ringkamp, 2009). Several molecular thermosensors have been identified in cutaneous peripheral nerve endings (Dhaka et al., 2006; Vriens et al., 2014). There is wide consensus that Transient Receptor Potential Melastatin 8 (TRPM8) channel, expressed in a restricted subpopulation of peripheral fibers (Dhaka et al., 2008; Takashima et al., 2007), is the main transducer for mild cold in somatosensory neurons (Bautista et al., 2007; Colburn et al., 2007; Dhaka et al., 2007; McKemy, 2018; McKemy et al., 2002; Peier et al., 2002). TRPM8 is also expressed at trigeminal epithelial endings innervating the tongue, cornea and upper airways (e.g. nose) (Alcalde et al., 2018; Dhaka et al., 2008; Takashima et al., 2007). Their activation by cold can trigger protective reflexes leading to tearing or rhinorrhea (Cruz & Togias, 2008; Parra et al., 2010; Quallo et al., 2015). Cold thermoreceptor endings also express various potassium and sodium channels that are important for tuning their temperature thresholds and peak firing rates (MacDonald et al., 2021; Madrid et al., 2009; Noel et al., 2009; Viana et al., 2002).

Molecular mechanism for noxious cold sensing have been more controversial (reviewed by Lewis & Griffith, 2024; Lolignier et al., 2016; MacDonald et al., 2020). Bioinformatic data mining identified Transient Receptor Potential Ankyrin 1 (TRPA1) which is activated by temperatures below 17 °C in recombinant expression systems (Story et al., 2003). In the DRG it was found in a subpopulation of neurons expressing nociceptive markers (e.g. TRPV1, CGRP) and non-overlapping with TRPM8. TRPA1 was proposed as a sensor for painful or noxious cold (Story et al., 2003). Subsequent studies found that TRPA1 is activated by a variety of endogenous and exogenous electrophilic compounds (Bandell et al., 2004; Jordt et al., 2004; reviewed by Zygmunt & Hogestatt, 2014). It is well accepted that TRPA1 plays a critical role as a molecular sentinel for chemical irritants and tissue damage in the skin and various internal organs (reviewed by Bautista et al., 2013; Talavera et al., 2020; Viana, 2016). The participation of TRPA1 in noxious cold sensing in trigeminal and DRG neurons has been validated in some studies (Bandell et al., 2004; Bernal et al., 2021; Gentry et al., 2010; Karashima et al., 2009; Kwan et al., 2006; Memon et al., 2017; Story et al., 2003) but not in others (Bautista et al., 2006; Jordt et al., 2004; Kwan et al., 2009; Nagata et al., 2005). Adding to this debate, the role of TRPM8 as a thermosensor has expanded beyond innocuous cold detection, with several studies suggesting an important role for TRPM8 in cold pain (Bautista et al., 2007; Colburn et al., 2007; Dhaka et al., 2007; Knowlton et al., 2010; Knowlton et al., 2013; Lippoldt et al., 2016; Pogorzala et al., 2013).

Visceral organs are also heavily innervated by functionally and molecularly diverse sensory endings, sending their peripheral projections to somatic and vagal ganglia (Prescott & Liberles, 2022). This innervation represents the sensory link between the viscera and the brain, critical for integrating information about the inner state of the body (Berntson & Khalsa, 2021; Chen et al., 2021). A fraction of vagal sensory neurons is activated by cold, but their mechanisms of cold sensitivity remain unclear. In addition, cold sensitivity in different vagal territories is currently unknown. Previous studies found low expression of TRPM8 in vagal ganglia (Hondoh et al., 2010; Kupari et al., 2019; Nassenstein et al., 2008). In contrast, vagal neurons show strong TRPA1 expression (Patil et al., 2023), and a role for TRPA1 in cold sensing in rat and mouse vagal neurons has been established before (Fajardo et al., 2008). The lungs and lower airways relay most of their sensory input through the vagus nerve (Berthoud & Neuhuber, 2000; Mazzone & Undem, 2016). Breathing cold air can trigger protective responses like cough, bronchoconstriction and mucosal secretion (Banner et al., 1985). It can also trigger asthma attacks in winter sport athletes (Larsson et al., 1993). A large fraction of airway-innervating vagal neurons express TRPA1 (Mazzone et al., 2020; Nassenstein et al., 2008; Patil et al., 2023) and one of their functions is to detect potentially harmful stimuli reaching the airways (Bessac et al., 2008; Mazzone & Undem, 2016).

Transduction mechanisms used by vagal sensory neurons are still poorly defined at the molecular level (Prescott & Liberles, 2022). To characterize the molecular mechanisms of temperature sensing in mouse vagal and trigeminal neurons, we performed a side-by-side comparison, using calcium imaging in combination with pharmacological and genetic tools. We found that TRPA1 is the main cold transducer in vagal neurons across the whole range of temperatures tested, whereas TRPM8 is indispensable in low-threshold trigeminal somatosensory neurons, with TRPA1 also contributing to noxious cold responses. In both ganglia we found a small fraction of neurons responding to cold by TRPA1- and TRPM8-independent mechanisms. Finally, we found that airway-innervating vagal neurons are particularly sensitive to temperature decreases by a mechanism engaging TRPA1.

## Results

### There are more cold-sensitive neurons in mouse VG than in TG

To compare the characteristics of cultured cold-sensitive (CS) visceral vagal (VG) and somatosensory trigeminal ganglion (TG) neurons, the bath temperature was rapidly lowered from ∼33 to ∼11 °C. As shown in figure 1A-B, CS neurons were found in both ganglia and their percentages were quantified in figure 1C. There were more CS neurons in VG than in TG (21.5 % (284/1321 VG cells) *vs* 14.6 % (154/1053 TG cells; p<0.001, Fisheŕs exact test). The distribution of response amplitudes to cold temperature was similar in both ganglia (figure 1D) with no difference in their means (0.52 ± 0.03 (VG) *vs* 0.58 ± 0.04 (TG), p=0.20, unpaired t-test). Most TRPM8-expressing neurons in mouse DRG have small somas (Dhaka et al., 2008). Measuring their cell diameters, we found that CS TG neurons were smaller in size on average (20.6 ± 0.3 µm *vs* 17.4 ± 0.3 µm, p<0.001, unpaired t-test), whereas there was no difference between the mean cell diameters of CS and cold-insensitive (CI) VG neurons (20.6 ± 0.3 µm *vs* 20.6 ± 0.2 µm, p=0.92, unpaired t-test, figure 1E). Figure 1F shows that cold responses in both ganglia covered the whole range of temperature thresholds tested. There was a clear increase in the number of high-threshold neurons (<19 °C) in VG, making the mean temperature threshold of VG CS neurons significantly lower compared to TG (19.2 ± 0.3 (VG) *vs* 20.7 ± 0.5 (TG) °C, p=0.006, unpaired t-test). These results indicate morphological differences between CS mouse vagal and trigeminal neurons, as well as functional differences in their overall response to cold temperature.

**Figure 1:**
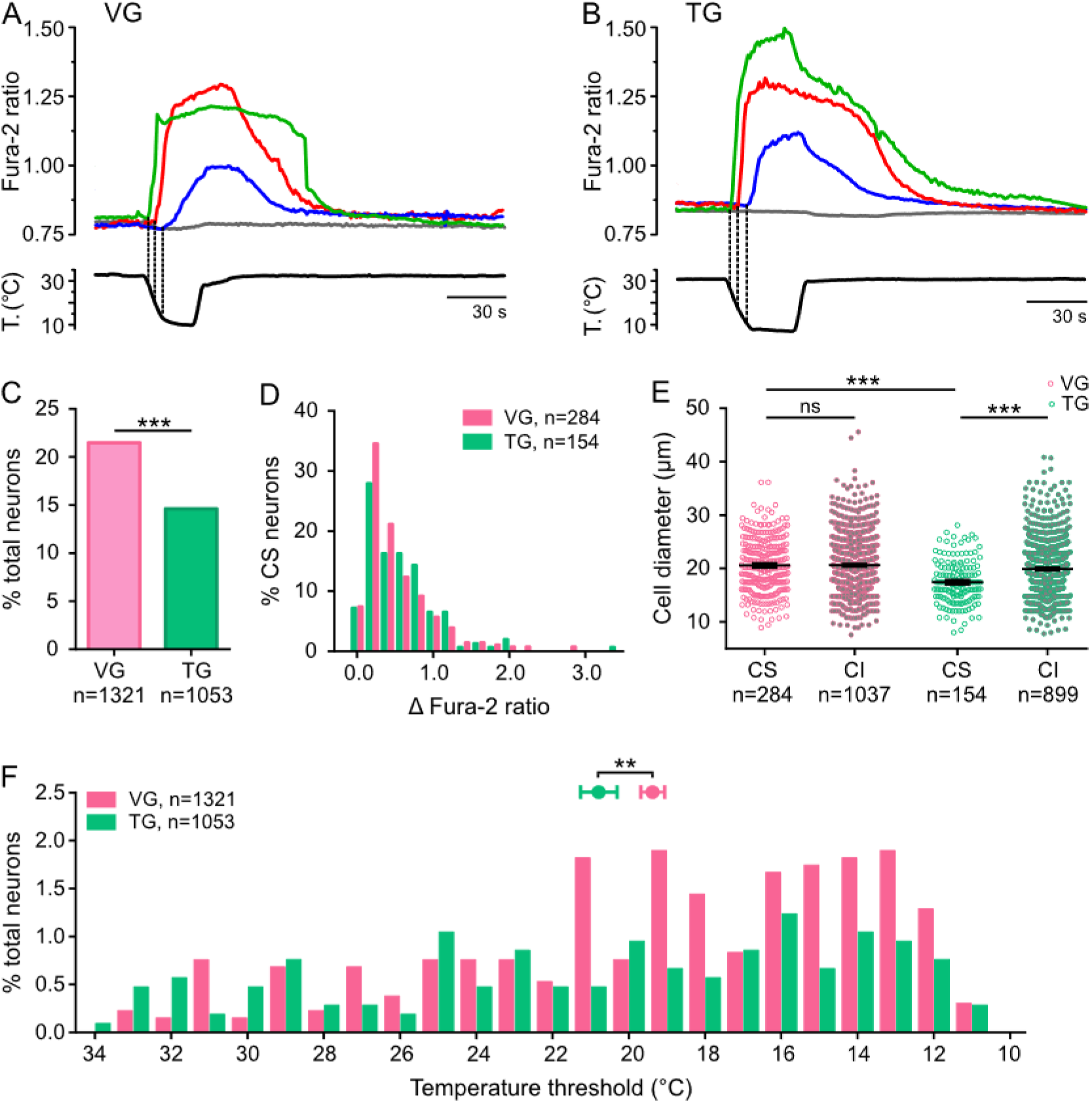
Differential characteristics of cold-sensitive (CS) neurons in VG and TG. Exemplary intracellular calcium traces of one cold-insensitive and three CS vagal (A) and trigeminal (B) neurons responding to different temperature thresholds. Neurons in each panel were recorded simultaneously. (A) Temperature thresholds (°C) are 29.3 (green), 17.5 (red) and 12.2 (blue). (B) Temperature thresholds (°C) are 29.5 (green), 22.6 (red) and 14.1 (blue). (C) Percentage of neurons responding to a decrease in temperature in VG and TG (p<0.001, Fisheŕs exact test). (D) Histogram showing the distribution of amplitude responses to a cold stimulus in VG and TG. (E) Cell diameters of cold-sensitive (CS) and cold-insensitive (CI) neurons (*** p<0.001, one-way ANOVA with Tukeýs post hoc). (F) Histogram showing temperature threshold distribution of CS neurons in VG and TG. Mean ± SEM for VG (pink, 19.2 ± 0.33) and TG (green, 20.7 ± 0.51) are indicated (** p<0.01, unpaired t-test).

### VG CS neurons are predominantly TRPA1-expressing, whereas the majority of TG CS neurons show a TRPM8-like phenotype

To determine whether these CS neurons express TRPA1 or TRPM8 channels, we exposed cultured VG and TG neurons to a cold ramp followed by the application of a second cold ramp combined with 100 µM menthol, and 100 µM AITC as shown in figure 2A,C. Since menthol can activate both TRPM8 and TRPA1 (Karashima et al., 2007; Lemon et al., 2019; McKemy et al., 2002; Peier et al., 2002; Xiao et al., 2008) but micromolar AITC is a selective TRPA1 agonist (Bandell et al., 2004; Jordt et al., 2004), we used the response to these agonists to classify CS neurons into three groups: TRPA1-like (AITC^+^, Menthol^+/-^), TRPM8-like (AITC^-^, Menthol^+^), and TRPA1-/TRPM8-independent (AITC^-^, Menthol^-^). Figure 2B,D clearly points to a different involvement of TRPA1 and TRPM8 in the cold sensing of VG and TG neurons. In VG, 68 % of CS neurons also responded to AITC, suggesting TRPA1 expression, whereas only 3 % showed a TRPM8-like phenotype. To verify the low TRPM8 expression in VG, we used a reporter transgenic mouse that expresses EYFP in TRPM8-positive neurons (Morenilla-Palao et al., 2014). In histological sections stained for GFP and tubulin β III, we found that TRPM8^+^ neurons represent only 4.5 % of all neurons in VG (figures 2H,I). Moreover, these few TRPM8-expressing neurons were predominantly located in the jugular ganglion (JG), at the rostral end of the VG complex (figures 2J,K). The low TRPM8 expression was further confirmed by quantitative PCR results from MACS purified VG neurons (see Methods, supplementary figure 1A).

**Figure 2:**
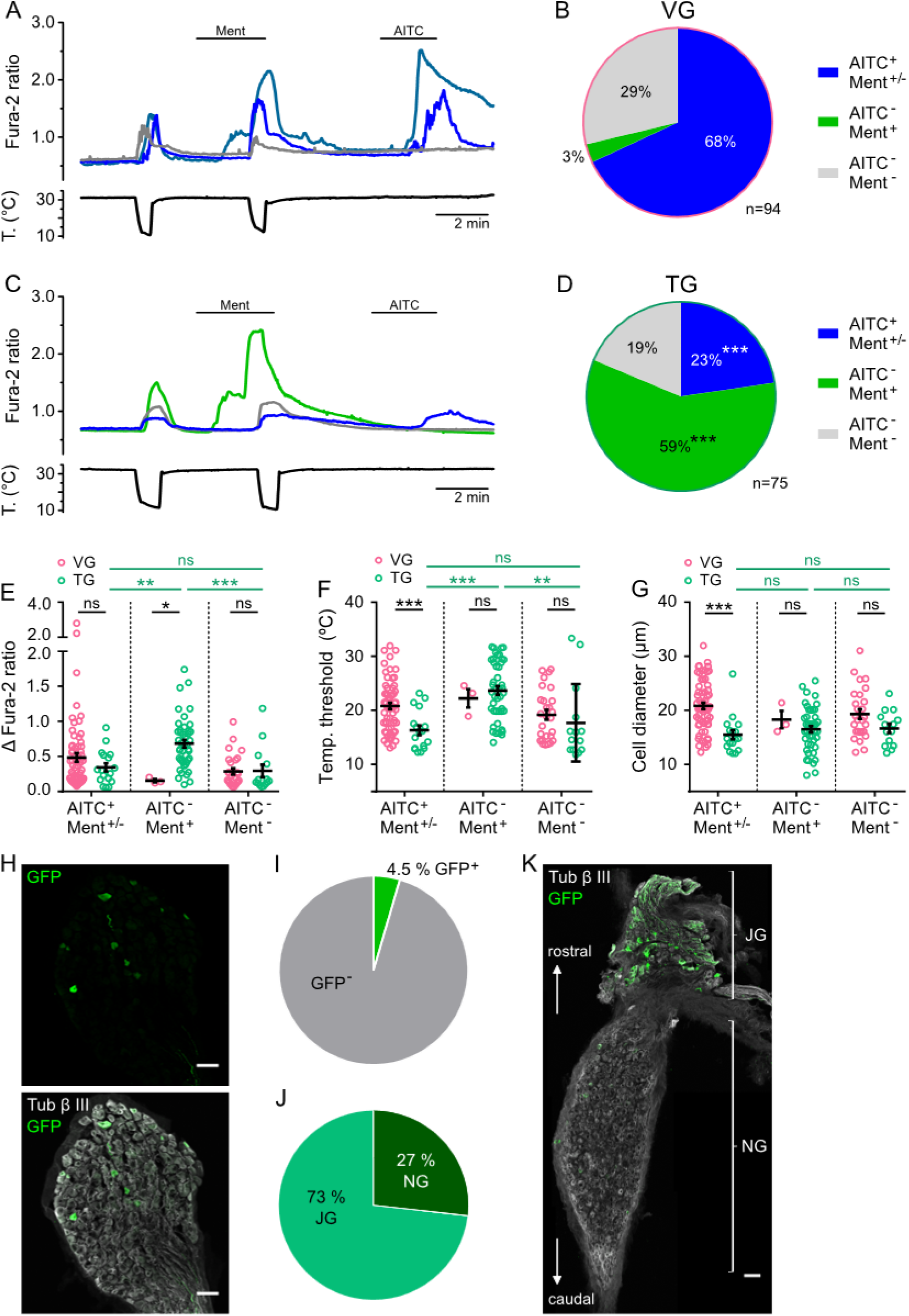
Different response profile to TRP agonists in cold-sensitive (CS) neurons in VG and TG. (A,C) Exemplary intracellular calcium traces for different classes of CS neurons: TRPA1-like (AITC^+^, Menthol^+/-^, blue trace), TRPM8-like (AITC^-^, Menthol^+^, green trace) and TRPA1-/TRPM8-independent (AITC^-^, Menthol^-^, grey trace) in VG (A) and TG (C) neurons, 100 µM menthol, 100 µM AITC. (B,D) Pie charts showing the percentage of CS cells split into different classes according to their responsiveness to menthol and AITC for VG (B) and TG (D) (*** p<0.001, Fisheŕs exact test, VG n=94, TG n=75 cells). Amplitudes (E), temperature thresholds (F), and cell diameters (G) of CS neurons split into the indicated groups according to their AITC and menthol sensitivity, one-way ANOVA with Sidak’s post hoc test, * p<0.05, ** p<0.01, *** p<0.001, VG AITC^+^ Ment^+/-^ n=64, VG AITC^-^ Ment^+^ n=3, VG AITC^-^ Ment^-^ n=27, TG AITC^+^ Ment^+/-^, n=17, TG AITC^-^ Ment^+^ n=44, TG AITC^-^ Ment^-^ n=14 cells. (H) Confocal image of a fixed and stained VG slice from a Trpm8^BAC-EYFP^ mouse, scale bar 50 µm. (I) Percentage of GFP^+^ neurons out of total beta tubulin^+^ neurons, n=3149 cells. (J) Pie chart showing the distribution of GFP^+^ neurons in either NG or JG, n=273 cells. (K) Mosaic image of a fixed and stained mouse VG, indicating NG and JG regions, scale bar 50 µm.

On the other hand, functional analysis of cold responses in the TG revealed that 59 % of CS neurons responded to menthol but not to AITC, characteristic of TRPM8 expression. Importantly, almost a quarter (23 %) of CS trigeminal neurons also responded to AITC (TRPA1-like). Of note, both ganglia contained a percentage of CS neurons that use a TRPA1-/TRPM8-independent cold transducing mechanisms (29 % in VG, 19 % in TG, figure 2B,D). The sensitivity to capsaicin in CS neurons was the same in both ganglia (31.9 vs 29.3 %, p=0.74, Fisheŕs exact test). In TG, AITC^-^ Menthol^+^ CS neurons showed larger amplitudes and lower temperature thresholds compared to the other two groups (figure 2E,F). This is in accordance with previous reports that TRPM8 is the main transducer for mild cold in somatic neurons (Bautista et al., 2007; Colburn et al., 2007; Dhaka et al., 2007). In contrast, in VG, there was no difference in the response amplitudes, temperature thresholds or cell diameter between the three groups of CS neurons (figure 2E-G). In addition, VG AITC^+^ CS neurons showed lower mean temperature thresholds and larger cell diameters compared to their TG counterparts (figure 2E-G).

### The majority of vagal cold temperature responses are mediated by TRPA1

To confirm TRPA1 as the main cold transducing molecule in vagal CS neurons, we used pharmacological and genetic approaches. As shown in figure 3A, we applied two cold ramps to VG neurons followed by an application of AITC. The first cold ramp was applied in control solution (black trace) or in the presence of a TRPA1 or TRPM8 antagonist. In the presence of two different TRPA1 antagonists, HC-030031 or A967079, many cold-evoked responses were abolished or strongly diminished (blue and red traces). In contrast, the majority of cold responses in vagal neurons were not affected in the presence of the potent TRPM8 antagonist RQ00203078 (green trace). These results are quantified in figure 3B. HC-030031 and A967079 reduced the number of CS neurons by 81 % and 46 %, respectively, compared to the control stimulus after the wash period. In radioligand binding assays, HC-030031 has been shown to lack full TRPA1 selectivity (Eid et al., 2008), which might explain why it blocks more CS responses than A967079. The response amplitudes of the remaining CS neurons were strongly reduced in the presence of these TRPA1 antagonists (figure 3C), and the large majority of inhibited neurons also responded to AITC (figure 3D), suggesting TRPA1 expression and selectivity of the two antagonists. In line with the low expression of TRPM8 we and others found in mouse vagal ganglia (Kupari et al., 2019), the TRPM8 antagonist RQ00203078 caused only a small decrease in the fraction of CS neurons (15.2 *vs* 19.7 %, p=0.29, Fisheŕs exact test, figure 3B). Furthermore, the mean amplitudes in neurons responding to cold in the presence of RQ00203078 or after a wash period were similar (0.31 ± 0.07 *vs* 0.32 ± 0.04, p=1.0, one-way ANOVA with Sidak’s post hoc test, figure 3C). Unexpectedly, 7 out of 11 neurons (64 %) that were inhibited by RQ00203078 were also AITC-sensitive (figure 3D).

**Figure 3:**
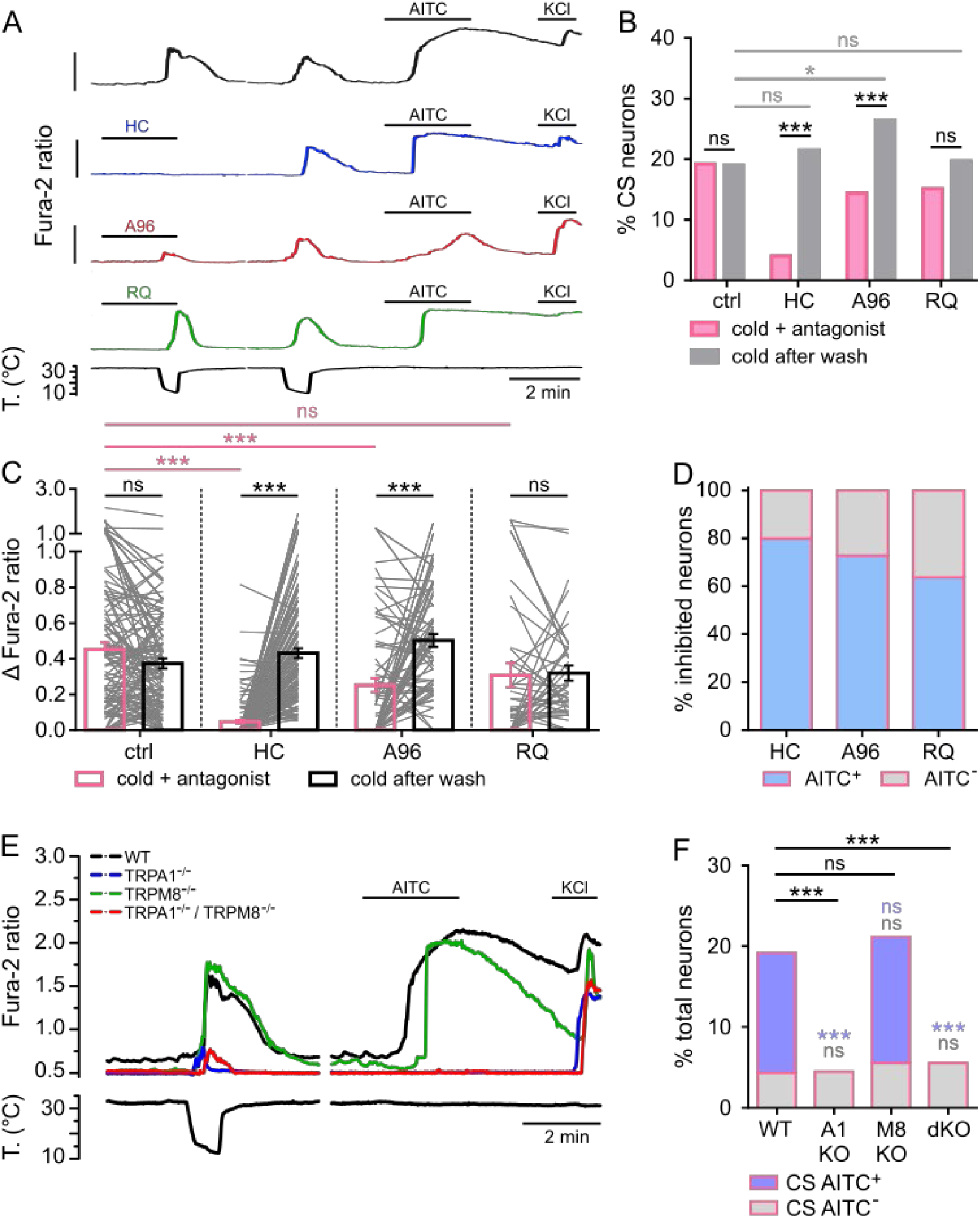
The majority of vagal cold responses are mediated by TRPA1. (A) Representative calcium traces of VG neurons from WT mice that were exposed to a first cold ramp in the presence of control solution (black trace) or antagonists (20 µM HC-030031 (HC, blue), 1 µM A967079 (A96, red) or 1 µM RQ00203078 (RQ, green)). A second cold ramp was applied after a wash period (3 min for control and HC, 6-7 min for A96 and RQ). Gap in x-axis (time) to align different traces with different lengths of wash periods, 100 µM AITC, 50 mM KCl. All calibrations represent a ΔF = 1, starting at 0.5. (B) Percentage of cold responders to the first cold stimulus (cold + control or antagonist, pink) and to the second cold stimulus (cold after wash, grey), Fisheŕs exact test, * p<0.05, *** p<0.001, ctrl n= 531, HC n=566, A96 n=257, RQ n=198 cells. (C) Mean amplitude of responses to the first cold stimulus (cold + control/antagonist, pink) and to the second cold stimulus (cold after wash, grey), light grey lines represent amplitudes of individual neurons, one-way ANOVA with Sidak’s post hoc, *** p<0.001, ctrl n= 113, HC n=122, A96 n=70, RQ n=41 cells. (D) Percentage of CS inhibited neurons that also responded to AITC (AITC^+^) or not (AITC^-^). Neurons that did not respond to cold in the presence of an antagonist but responded to cold after a wash period were counted as inhibited neurons, HC n=99, A96 n=33, RQ n=11 cells. (E) Exemplary traces of CS neurons from mice with the indicated genotypes. Gap in x-axis (time) to align different traces from different experiments, a second cold stimulus was applied before AITC application (not shown here),100 µM AITC, 50 mM KCl. (F) Percentage of vagal neurons that responded to a decrease in temperature in mice with different genotypes, Fisheŕs exact test compared to WT, *** p<0.001. The total number of cells recorded: WT n=531, A1 KO n=245, M8 KO n=217, dKO n=289 cells, AITC sensitivity is indicated in blue.

To investigate whether we missed some of the CS neurons due to insufficient washing out of the antagonist, we performed additional experiments with a modified protocol. We applied three cold ramps: cold in control conditions, cold in the presence of RQ00203078, and cold after a 6-7 minute washout (supplementary figure 2A). The number of VG neurons responding to a first cold ramp (control) were similar to the ones responding to the third cold ramp (after washout) (21.5 % *vs* 19.8 %, p=0.87, Fisheŕs exact test, supplementary figure 3B), showing that we did not miss any CS neurons in our previous protocol due to insufficient washout of the antagonist. One of the few CS vagal neurons blocked by the TRPM8 antagonist RQ00203078 is shown (blue trace). Again, 6 out of 8 neurons (75 %) inhibited by RQ00203078 responded to AITC (supplementary figure 3D), suggesting that RQ00203078 has some inhibitory effect on vagal CS TRPA1-expressing neurons. This may indicate some co-expression of TRPM8 and TRPA1 in VG, unlike results obtained in DRG (Story et al., 2003).

Next, we compared cold and AITC sensitivity in vagal neurons from mice with different genotypes: wild type, Trpa1^-/-^, Trpm8^-/-^, or Trpa1^-/-^::Trpm8^-/-^ (dKO) (figures 3E-F). The results clearly show that the frequency of responses to a cold stimulus were strongly reduced in nodose cultures from Trpa1^-/-^ or Trpa1^-/-^::Trpm8^-/-^ mice (19.2 % WT *vs* 4.5 % Trpa1 KO, p<0.001, 19.2 % WT *vs* 5.5 % dKO, p<0.001, Fisheŕs exact test). In mice lacking TRPA1, the few remaining CS neurons did not respond to AITC (figures 3E-F). Furthermore, these remaining CS neurons displayed similar characteristics to AITC^-^ CS neurons in cultures from WT or TRPM8^-/-^ mice: smaller response amplitudes, higher temperature thresholds and smaller cell diameters (supplementary figures 3A-C).

In summary, using agonists, antagonists, and neuronal cultures from knock out mice for the different cold transducers, the data show that vagal cold responses are mainly mediated via TRPA1 with minimal involvement of TRPM8. These AITC^+^ neurons span the entire range from low to very high threshold CS neurons. A smaller percentage of vagal CS neurons harbor a molecular mechanism independent of TRPA1 or TRPM8; these neurons show smaller response amplitudes and cell diameter as well as higher temperature thresholds. The molecular identity of this cold sensor remains to be determined.

### TRPM8 and TRPA1 show differential contributions to trigeminal cold responses

The same experimental strategy used in VG was applied to TG cultures to investigate their cold sensitivity (figure 4A). The involvement of TRPA1 and TRPM8 in cold sensing was very different in TG compared to VG. In control conditions (i.e. without antagonist), 15.5 % of TG neurons responded to a first cold ramp (figure 4B). Application of a cold ramp in the presence of the TRPM8 antagonist RQ00203078 reduced the fraction of CS neurons to 5.2 %. Calculated from responses obtained after washing out the drug, this represents a decrease of 55 %. This is an underestimate of TRPM8-dependent responses considering that RQ wash was incomplete (see below). In contrast, the fraction of CS neurons in the presence of the selective TRPA1 antagonist A967079 was similar to control conditions (17.5 % (A96) *vs* 15.5 % (ctrl), p=0.67, Fisheŕs exact test) and to the fraction of CS neurons after having washed out the compound (17.5 % *vs* 25.0 %, p=0.13, Fisheŕs exact test). In the case of the TRPA1 antagonist HC-030031, 49 % of TG CS neurons were blocked when compared to neurons responding to cold after washout (9.6 % vs 18.9 %, p=0.003, Fisheŕs exact test, figure 4B), however, this antagonist has been shown to be not fully selective for TRPA1 (Eid et al., 2008). Interestingly, when comparing the responses to the first cold stimulus between neurons that were exposed to one of the antagonists and neurons in control conditions, we found that all three antagonists significantly decreased the mean amplitudes of the responses (figure 4C). This suggests the involvement of both channels, TRPM8 and TRPA1, in cold responses of TG neurons. We also compared the amplitudes of the responses to the second cold ramp. Compared to the control group, the mean amplitudes were smaller after the wash period following A967079 (0.43 ± 0.06 (ctrl) *vs* 0.23 ± 0.02 (A96), p=0.02, one-way ANOVA with Sidak’s post hoc test) and RQ00203078 (0.43 ± 0.06 (ctrl) *vs* 0.18 ± 0.03 (RQ), p=0.03, one-way ANOVA with Sidak’s post hoc test) application. Furthermore, when looking at the inhibited neurons, i.e. those that responded to a cold stimulus after washout (control) but did not respond in the presence of the antagonist, we found that 9 out of 12 (75 %) neurons that were inhibited by RQ00203078 also responded to AITC (figure 4D), which is not characteristic for TRPM8-expressing neurons. These results suggested that our protocol might not allow sufficient washout times. Therefore, we changed the protocol, with the application of RQ sandwiched between a control and a wash cold ramp (supplementary figure 4A). In the presence of RQ, the percentage of CS neurons was reduced from 16.0 % to 8.3 % (p = 0.003, Fisheŕs exact test). Moreover, we found a significant decrease in the number of CS neurons (16.0 % *vs* 10.4 %, p=0.04, Fisheŕs exact test, supplementary figure 4B) and a clear reduction in the response amplitudes (0.67 ± 0.10 *vs* 0.26 ± 0.03, p<0.001, one-way ANOVA with Sidak’s post hoc test, supplementary figure 4C) when comparing the cold responses to the first (control) and the third (after a 6-7 minute wash) cold stimuli. This strongly suggests the incomplete washout of RQ and indicates that we probably missed a fraction of TG CS neurons in our previous protocol (figures 4A-D). When looking at the AITC sensitivity of the inhibited neurons using the protocol with three cold ramps, we found that 26 out of 33 neurons (78.8 %) that were inhibited by RQ00203078 did not respond to AITC, showing a TRPM8 phenotype (supplementary figure 5D).

**Figure 4:**
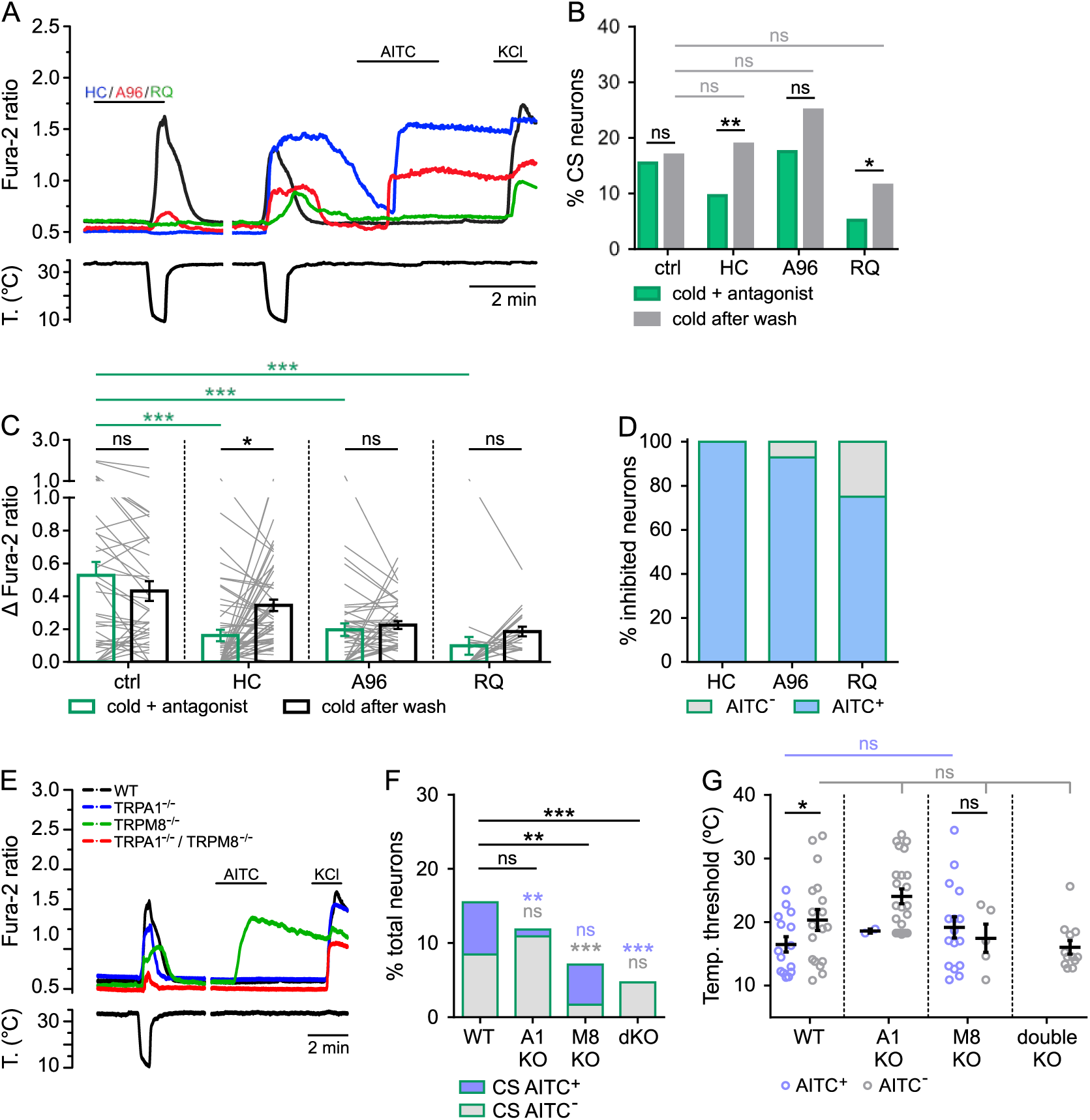
TRPA1 and TRPM8 are involved in cold sensing in TG neurons. (A) Representative calcium traces of TG neurons from WT mice that were exposed to a first cold ramp in the presence of control solution (black trace) or antagonists (20 µM HC-030031 (HC, blue), 1 µM A967079 (A96, red) or 1 µM RQ00203078 (RQ, green)). A second cold ramp was applied after a period of wash (3 min for control and HC, 6-7 min for A96 and RQ). Gap in x-axis (time) to align different traces with different lengths of wash periods, 100 µM AITC, 50 mM KCl. (B) Percentage of cold responders to the first cold stimulus (cold + antagonist, green) and to the second cold stimulus (cold after wash, grey), Fisheŕs exact test, * p<0.05, ** p<0.01, control n= 213, HC n=270, A96 n=160, RQ n=174 cells. (C) Mean amplitudes to the first cold stimulus (cold + antagonist, green) and to the second cold stimulus (cold after wash, grey), light grey lines represent amplitudes of individual neurons, one-way ANOVA with Sidak’s post hoc, * p<0.05, *** p<0.001, ctrl n= 38, HC n=52, A96 n=42, RQ n=21 cells. (D) Number of inhibited neurons that also responded to AITC (AITC^+^) or not (AITC^-^). Neurons that did not respond to cold in the presence of an antagonist but responded to cold after a wash period were counted as inhibited neurons, HC n=26, A96 n=14, RQ n=12 cells. (E) Exemplary traces of CS neurons from mice with the indicated genotypes. Gap in x-axis (time) to align different traces from different experiments, a second cold stimulus was applied before AITC application (not shown here), 100 µM AITC, 50 mM KCl. (F) Number of trigeminal neurons, out of total, that responded to a decrease in temperature in mice with different genotypes, Fisheŕs exact test compared to WT, ** p<0.01, *** p<0.001, WT n=213, A1 KO n=211, M8 KO n=296, dKO n=255 cells, AITC sensitivity is indicated in blue. Temperature thresholds of cold responses (G), amplitudes (H) and cell diameters (I) for CS TG neurons from different genotypes, blue circles represent AITC^+^ neurons, grey circles indicate AITC^-^ neurons, one-way ANOVA with Tukeýs post hoc, * p<0.05, WT AITC^+^ n=15, WT AITC^-^ n=18, A1 KO AITC^+^ n=2, A1 KO AITC^-^ n=23, M8 KO AITC^+^ n=16, M8 KO AITC^-^ n=5, dKO AITC^+^ n=0, dKO AITC^-^ n=12 cells.

Pharmacological tools are seldom fully selective. Thus, we used TG neurons from transgenic mice lacking specific temperature transducers and tested their cold sensitivity (figure 4E). In this experimental series, 15.5 % of wild type TG neurons were CS and, of those, 45 % responded to AITC (figure 4F). CS AITC^+^ neurons were virtually abolished in Trpa1^-/-^ mice (7.0 *vs* 0.9 %, p=0.002, Fisheŕs exact test). However, this abolishment did not cause a significant reduction in the total number of CS neurons (15.5 % WT *vs* 11.8 % A1 KO, p=0.32, Fisheŕs exact test). On the other hand, a significant reduction was found in Trpm8^-/-^ mice (15.5 % WT *vs* 7.1 % M8 KO, p=0.003, Fisheŕs exact test, figure 4F), which causes a 54 % reduction of CS neurons in Trpm8 KO mice compared to wild type mice, a similar reduction to that obtained with RQ. As expected, this reduction was found in the AITC^-^ population (8.5 *vs* 1.7 %, p<0.001, Fisheŕs exact test). Finally, in cultures obtained from double KO mice, the number of CS neurons was decreased further to 4.7 % which represents ∼30 % of CS neurons found in wild type animals. None of these CS neurons was AITC-sensitive. Similarly to the results obtained in vagal neurons, these results suggest the presence of a cold sensor independent of TRPM8 or TRPA1 in trigeminal neurons.

As shown previously, WT AITC^-^ CS neurons (potentially TRPM8^+^) had higher threshold temperatures than AITC^+^ (potentially TRPA1^+^) CS neurons (20.3 ± 1.7 *vs* 16.5 ± 1.2 °C, p=0.04, unpaired t-test, figure 4G). In alignment with the fact that TG TRPM8^+^ neurons have lower temperature thresholds, the population of low threshold AITC^-^ neurons was absent in neurons from Trpm8 KO mice (figure 4G). In neurons from Trpa1 KO mice there was not a single CS neuron responding to temperatures below 18 °C (figure 4G), indicating indeed some contribution of TRPA1 to noxious cold sensing in TG.

In summary, and taken together all results so far, TRPM8 emerges as the key cold transducer molecule in TG and responsible for mediating cold responses that are mainly defined as having low temperature thresholds. Furthermore, TRPA1 contributes to cold sensitivity at very low temperatures in TG to some extent, although there is no significant effect on the fraction of CS neurons when either blocking TRPA1 pharmacologically or abolishing the channel genetically. Notably, about 25 % of TG CS neurons engage a transduction mechanism alternative to TRPM8 and TRPA1.

### The potassium channel blocker 4-AP recruits CS neurons independently of TRPA1 and TRPM8 expression

Blockers of voltage gated potassium channels of the Kv1 family (e.g. micromolar 4-Aminopyridine (4-AP) or kM-conotoxin RIIIJ) recruit a population of cold-sensitive TG and DRG neurons that are cold-insensitive in their absence (Memon et al., 2017; Viana et al., 2002). In addition, several Kv1 channel blockers, including 4-AP, shift the temperature threshold of original CS neurons to higher temperatures, demonstrating its role as a molecular brake of thermal responses (Gonzalez et al., 2017; Madrid et al., 2009). Furthermore, a reduction in this molecular brake plays a major role in cold hypersensitivity following nerve injury (Piña et al., 2019). In qPCR experiments from RNA extracted from MACS purified neurons (see Methods) we found very similar transcript levels for Kv1.1, the main component of the brake current (Madrid et al., 2009), in both ganglia (supplementary figure 1D). Next, we investigated the effect of 4-AP on cold sensitivity in VG and TG neurons. For this, we applied a cold stimulus in control conditions, followed by a second cold stimulus in the presence of 100 μM 4-AP. To examine the role of the brake current in different neuronal subpopulations, this protocol was also used in cultures of the different genotypes. In some experiments, selective TRPM8 (i.e. WS-12) and TRPA1 (i.e. AITC) agonists were applied after the two cold ramps (figures 5A-B). Using this protocol, we could characterize the population of recruited neurons and their expression of TRPA1 or TRPM8.

**Figure 5.**
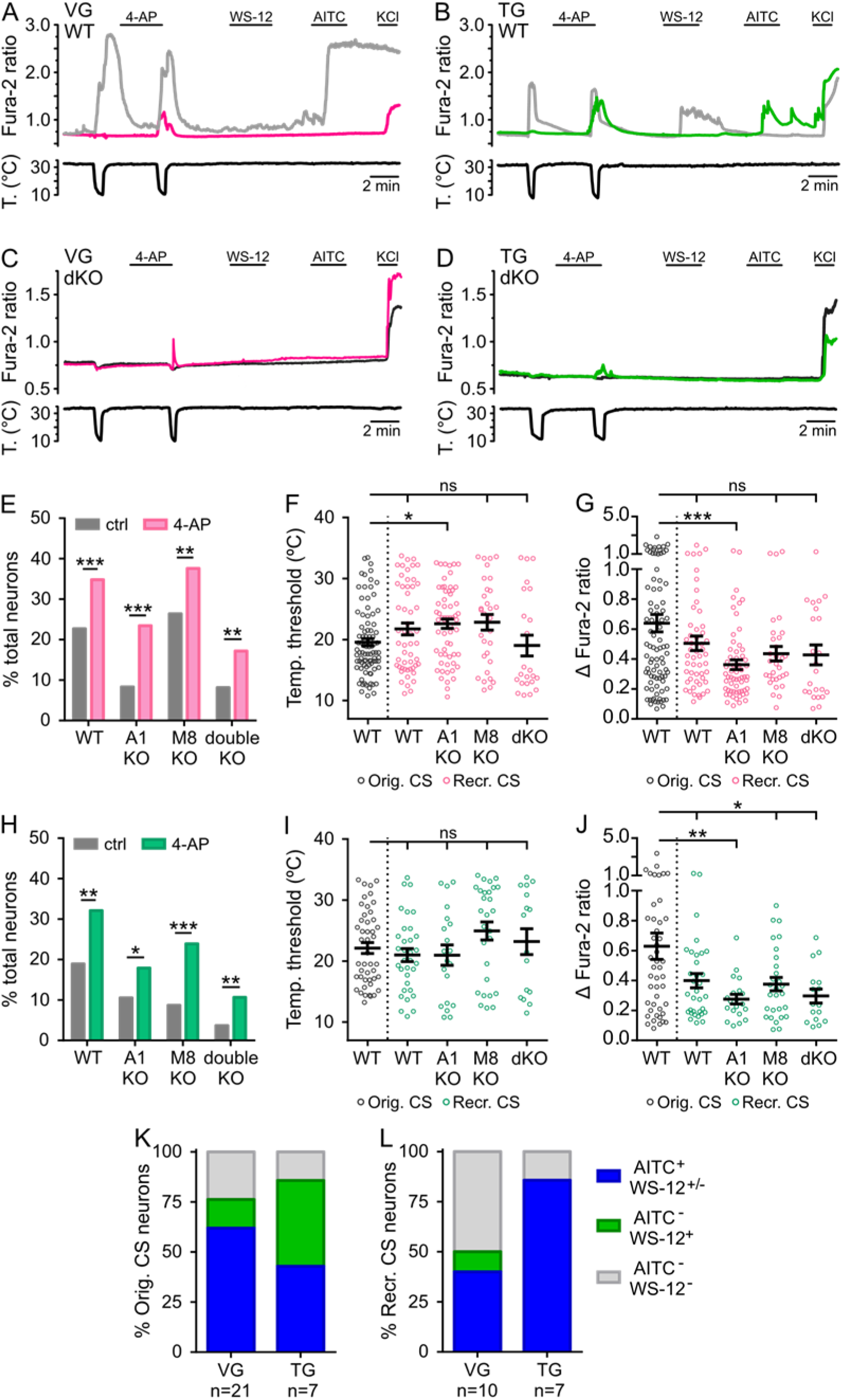
4-AP recruits CS neurons independently of TRPA1 and TRPM8 expression. Exemplary traces of original CS (grey), 4-AP recruited CS (pink/green) and CI (black) neurons from WT (A, B) and double KO (C, D) mice, 100 µM 4-AP, 500 nM WS-12, 100 µM AITC, 50 mM KCl. (E) Percentage of VG neurons that respond to a decrease in temperature in control conditions or in the presence of 100 µM 4-AP from mice with different genotypes, Fisheŕs exact test, ** p<0.01, *** p<0.001, WT n = 388, A1 KO n = 397, M8 KO n = 250, dKO n = 233 cells. Temperature thresholds (F) and amplitudes (G) of cold-evoked responses for recruited CS VG neurons from different genotypes (pink circles) and for original CS neurons from WT mice (grey circles), one-way ANOVA with Sidak’s post hoc, * p<0.05, *** p<0.001, original WT n = 88, recruited WT n = 53, A1 KO n = 61, M8 KO n = 31, dKO n = 23 cells. (H) Percentage of TG neurons that respond to a decrease in temperature in control conditions or in the presence of 100 µM 4-AP from mice with different genotypes, Fisheŕs exact test, * p<0.05, ** p<0.01, *** p<0.001, TG: WT n = 243, A1 KO n = 257, M8 KO n = 184, dKO n = 216 cells. Temperature thresholds (I) and amplitudes (J) of cold-evoked responses for recruited CS TG neurons from different genotypes (green circles) and for original CS neurons from WT mice (grey circles), one-way ANOVA with Sidak’s post hoc, * p<0.05, ** p<0.01, original WT n = 46, recruited WT n = 33, A1 KO n = 20, M8 KO n = 28, dKO n = 16 cells. (K, L) Stacked histograms showing the percentage of original (K) (VG n=21, TG n=7), or recruited (L) (VG n=10, TG n=7), CS neurons in WT mice that also responded to the different combinations of 100 µM AITC and 500 nM WS-12.

We found that 4-AP recruited a similar fraction of neurons previously insensitive to cold in VG (figure 5E) and TG (figure 5H), making up 13 % of the total number of cells in both ganglia (supplementary figures 5E,F). The sensitivity to WS-12 and AITC of recruited neurons revealed important differences in VG and TG. In VG, 50 % of recruited neurons were insensitive to AITC or WS-12, suggesting a TRPA1/TRPM8-independent mechanism, and only 1 out of 10 had a typical TRPM8-like phenotype (AITC^-^/WS-12^+^), consistent with the low expression of TRPM8 in VG. In contrast, in TG, 6 out of 7 recruited cells were AITC^+^ and none of them responded exclusively to WS-12 (figure 5L). This suggests that a pulse to 11 °C activates all TRPM8^+^ neurons in TG and that 4-AP recruits neurons from a different, mainly TRPA1-expressing subpopulation.

As clearly shown in figure 5E, in VG, the number of CS neurons in the presence of 4-AP was not only increased significantly in cultures from WT mice but also in cultures from Trpa1 KO, Trpm8 KO and double KO mice. Figure 5C shows one of these recruited VG neurons from a double KO mouse. 4-AP recruited CS VG neurons had variable temperature thresholds. In neurons from Trpm8 KO and double KO mice, their mean temperature threshold was similar to original CS neurons from WT mice (figure 5F). In neurons from Trpa1 KO mice, the mean temperature threshold was slightly but significantly increased. However, recruited CS neurons from all genotypes were found to span the entire temperature range (33 - 11 °C, figure 5F). The mean amplitudes of these recruited CS neurons were slightly smaller, although only with a statistically significant difference in the Trpa1 KO group (figure 5G). Similarly, 4-AP also increased the fraction of CS neurons in cultures from all four genotypes in TG neurons (figure 5H). Similar to VG, TG recruited neurons from all genotypes spanned the entire temperature range (figure 5I). As shown in figure 5J, the mean response amplitudes in recruited neurons were smaller than those of original CS neurons. A typical example of a recruited CS neuron in TG from a double KO mouse is shown in Figure 5D.

In addition to recruiting more neurons during a cold stimulus, we found that 4-AP shifted the temperature thresholds of original CS neurons to higher temperatures. For example, in VG from wild type mice the mean threshold temperature shifted from 19.7 to 24.1 °C (p < 0.001), while in TG it averaged 21.8 °C in control and 26.7 °C in the presence of 4.AP (p < 0.001). This effect was highly significant in neurons from all genotypes in both, VG and TG (supplementary figure 5A,C). In contrast, the amplitude of cold-evoked responses was not affected by 4-AP (supplementary figure 5B,C).

In summary, our results demonstrate that 4-AP recruits CS neurons in VG and in TG in animals lacking specific temperature sensors (TRPA1, TRPM8 or both). The responses in these recruited neurons are predominantly small in amplitude, their threshold is not restricted to a certain temperature range, and they are not responsive to the selective TRPM8 agonist WS-12. This suggests that 4-AP recruits neurons from different cold-sensitive populations.

### High expression levels of TRPA1 in vagal neurons

The functional, histochemical and qPCR results indicated a lower expression of TRPM8 in VG compared to TG. We posed the same question for TRPA1 expression. As shown in figure 6A, both ganglia had a similar percentage of AITC-responsive neurons (25.4 *vs* 25.1 %, p=0.94, Fisher’s exact test), suggesting similar number of TRPA1-expressing neurons. However, as shown in figure 6B, only a small fraction of AITC^+^ neurons responded to cold in TG. Remarkably, this fraction was much larger in VG than in TG (62.7 % *vs* 12.7 %, p<0.001, Fisher’s exact test). Moreover, in both ganglia, CS AITC^+^ neurons showed larger mean response amplitudes to AITC compared to cold-insensitive (CI) ones (figure 6C,D), suggesting that TRPA1 expression levels differ in VG and TG, and influence the cold sensitivity of TRPA1-expressing neurons. To test this hypothesis, we used magnetic-activated cell sorting (see Methods) to purify neurons from TG and VG and quantify their TRPA1 mRNA expression by qPCR. Consistent with the hypothesis, we found significantly higher TRPA1 mRNA expression levels in the latter (0.00099 *vs* 0.00056 2^ΔCt^, p=0.04, unpaired t-test, supplementary figure 1B). Additionally, the calcium imaging data shows that the mean calcium influx in response to AITC was larger in VG than in TG (0.83 ± 0.07 *vs* 0.52 ± 0.04, p<0.001, one-way ANOVA with Sidak’s post hoc test, figure 6E), a functional confirmation of the hypothesis.

**Figure 6.**
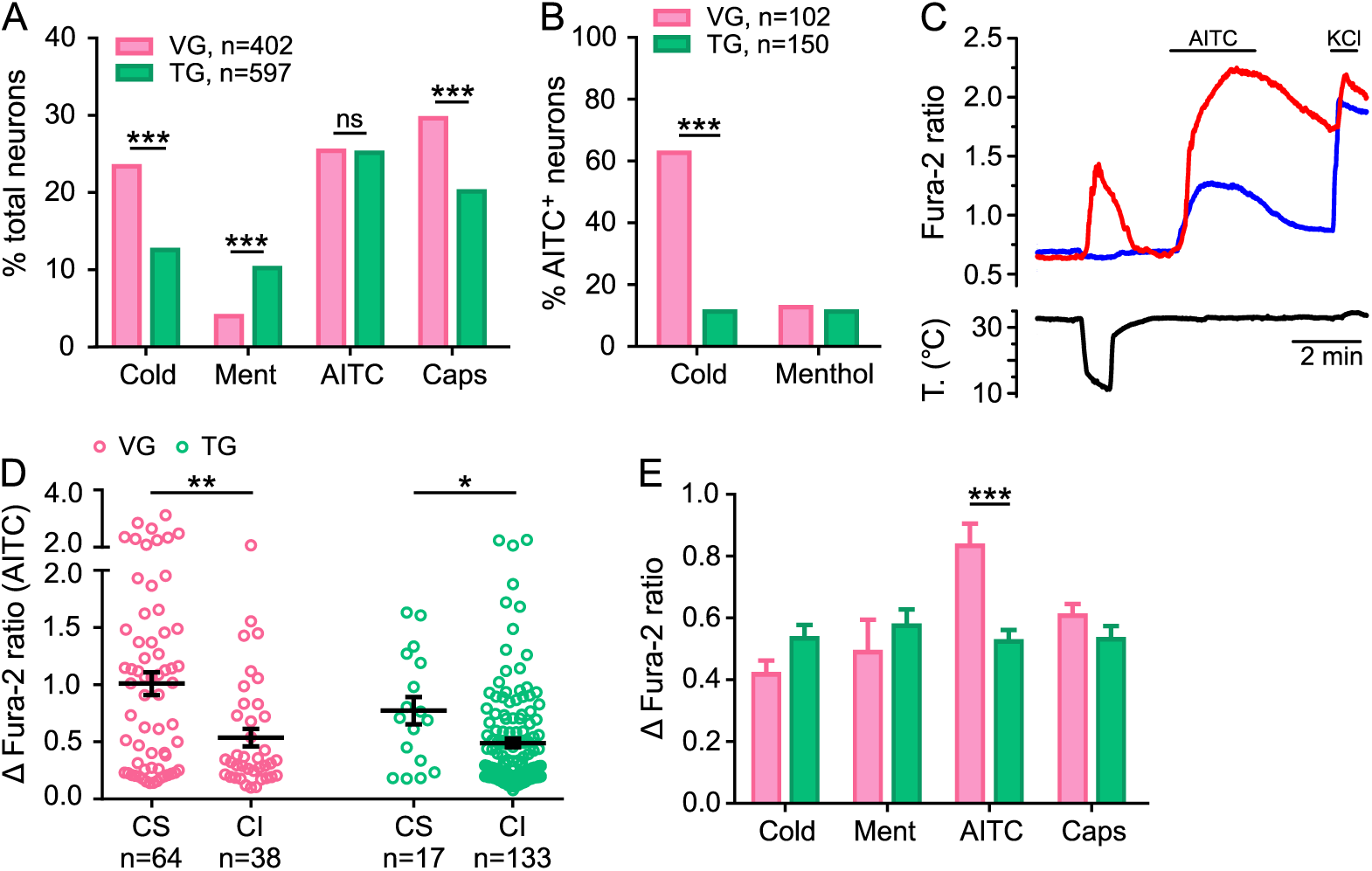
Vagal TRPA1-expressing CS neurons are more sensitive to TRPA1 agonists. (A) Percentage of total number of neurons responding to each stimulus in VG (pink) and TG (green), *** p<0.001, Fisheŕs exact test. (B) Number of cells responding to cold or menthol out of all AITC-sensitive neurons, *** p<0.001, Fisheŕs exact test. (C) Representative traces of a cold-sensitive (red) and a cold-insensitive (blue) vagal neuron. Both neurons are AITC^+^, 100 µM AITC, 50 mM KCl. (D) Amplitudes of response to AITC in cold-sensitive (CS) and cold-insensitive (CI) AITC^+^ neurons, * p<0.05, ** p<0.01, unpaired t-test. (E) Response amplitude to different stimuli in VG and TG neurons, *** p<0.001 one-way ANOVA with Sidak’s post hoc test.

Keeping in mind that both ganglia presented a similar number of TRPA1-expressing neurons overall, these data show that VG TRPA1-expressing neurons are more sensitive to TRPA1 agonists than trigeminal TRPA1-expressing neurons, probably by expressing more functional channels per neuron.

### There are more CS neurons in airway-innervating vagal neurons

The vagus nerve innervates different internal organs and tissues, including the gut, the heart and the lungs (Mazzone & Undem, 2016). Compared to other organs (e.g. the heart), the airways and lungs are one tissue innervated by the vagus nerve that are constantly exposed to temperature fluctuations during breathing, and inhaling cold air triggers powerful bronchomotor responses (Jammes et al., 1983). We asked whether VG neurons innervating the airways are particularly cold-sensitive. For this, mice were briefly anesthetized and intubated to add a small volume of retrograde tracer dye (DiI or WGA-Alexa 594) into the lungs (Kaelberer & Jordt, 2016), (see Methods). Immunohistochemical staining of paraformaldehyde-fixed tissues was performed from mice that were previously intubated with WGA-Alexa 594. Labelling was clearly present in a small fraction of neurons in sections from vagal ganglia (111/2998, 3.7 %, supplementary figure 6A), whereas no labelling was present in sections from lumbar DRG (0/1155, supplementary figure 6B). In addition, widespread labelling was observed in sections from the trachea and the lung parenchyma (supplementary figure 6C-D). Similar results were obtained from cultured and fixed neurons after retrograde labelling with the fluorescent marker DiI. The DiI crystals were seen as intracellular fluorescent dots within the soma of airway-innervating neurons (supplementary figure 7A). Following DiI application, 4.4 % (23/519) of VG cells were found to be DiI^+^, whereas cultures from TG or different sections of DRG only showed very few or no DiI^+^ cells (supplementary figure 7B).

Figure 7A shows calcium imaging records of one exemplary DiI^+^ (red trace) and two DiI^-^ (grey and black traces) vagal neurons during exposure to a cold ramp and a battery of agonists for different thermosensitive TRP channels. In these experiments, the more potent and selective TRPM8 agonist WS-12 was used instead of menthol (Bodding et al., 2007). As shown in figure 7B, more cold-sensitive neurons were found in the DiI^+^ population (35.8 *vs* 24.9%, p=0.03, Fisheŕs exact test). In addition, the percentage of AITC-sensitive neurons was also higher in the DiI^+^ population (41.1 *vs* 28.1%, p=0.01, Fisheŕs exact test). In contrast, there was no difference in the WS-12^+^ population (7.4 *vs* 6.1 %, p=0.64, Fisheŕs exact test). The number of capsaicin sensitive neurons was smaller in DiI^+^ cells than in DiI^-^ cells (49.5 *vs* 37.9 %, p=0.04, Fisheŕs exact test). In addition to the increased number of cold- and AITC-sensitive neurons overall, there was an increase in the number of neurons responding to cold out of the AITC^+^ population in DiI^+^ neurons (82.1 *vs* 60.4 %, p=0.013, Fisheŕs exact test, figure 7C). In accordance with this, the percentage of AITC^+^ CS neurons was significantly larger in DiI^+^ neurons compared to DiI^-^ neurons (33.7 *vs* 17.0 %, p<0.001, Fisheŕs exact test, figure 7D). We showed in figure 2B that 68.1 % of vagal CS neurons also responded to AITC. In VG neurons innervating the lungs and lower airways, this percentage was even higher, increasing to 94.1 % (p=0.002, Fisheŕs exact test). Not only did we find an increase in the number of neurons responding to cold and AITC, but their response amplitudes were also increased in DiI^+^ neurons as shown in figure 7E). Furthermore, there was an increase in very high threshold (< 19 °C) CS neurons in airway-innervating neurons compared to VG neurons innervating other tissues (figures 7F,G). These data show that, in vagal neurons innervating the lower airways, the responses to a decrease in temperature are frequent and mediated almost exclusively by TRPA1 channels.

**Figure 7.**
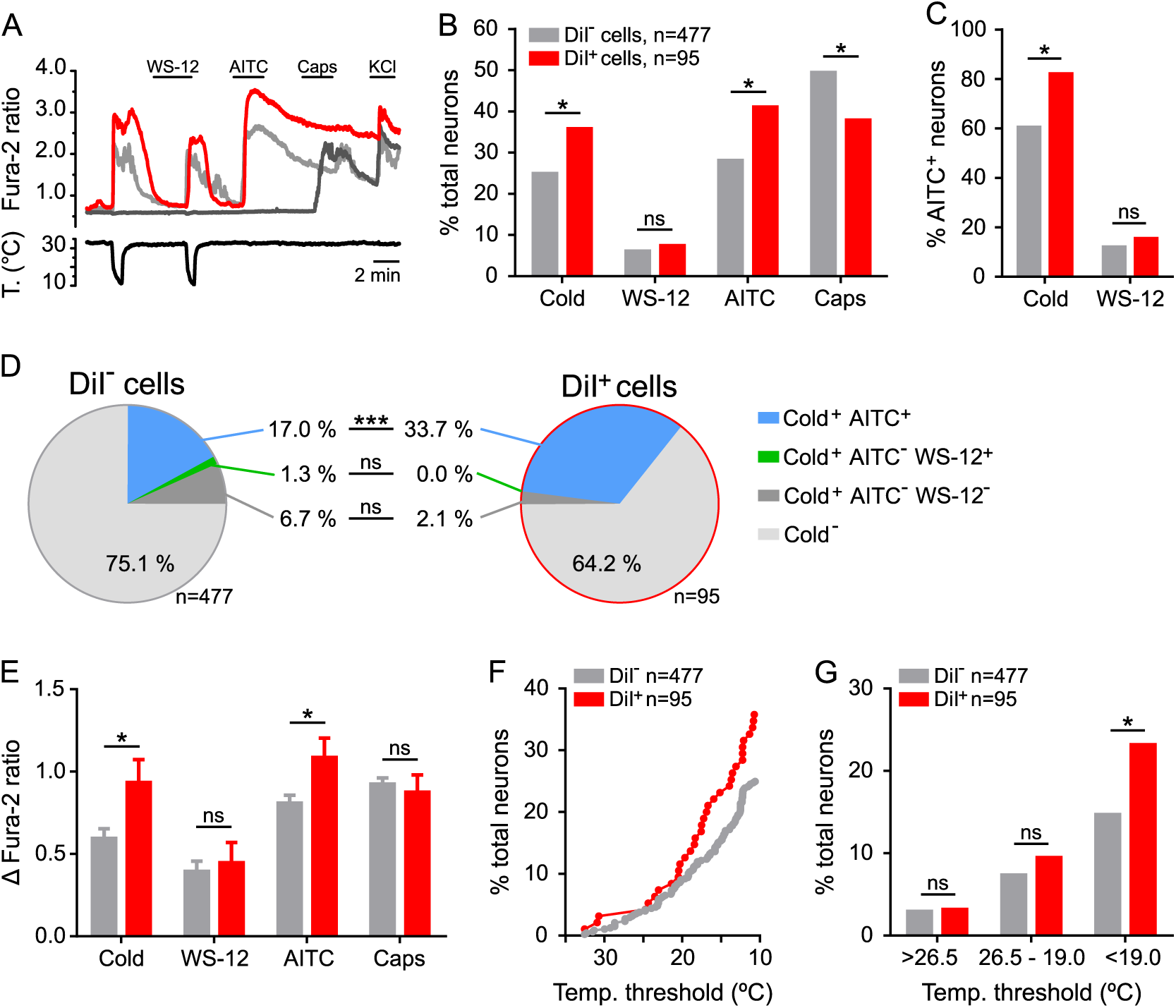
Airway-innervating vagal neurons are more cold- and AITC-sensitive. (A) Exemplary traces of one DiI^+^ (red) and two DiI^-^ (grey) neurons responding to different stimuli combinations, 500 nM WS-12, 100 µM AITC, 100 nM Capsaicin, 50 mM KCl. (B) Percentage of total number of DiI^+^ and DiI^-^ neurons responding to each stimulus, * p<0.05 Fisheŕs exact test. (C) Number of cells responding to cold or WS-12 out of all AITC-sensitive neurons, * p<0.05 Fisheŕs exact test, DiI^-^ n=134, DiI^+^ n=39 cells. (D) Pie charts showing the percentage of total cells that respond to cold and AITC (Cold^+^ AITC^+^), WS-12 but not AITC (Cold^+^ AITC^-^ WS-12^+^), or neither AITC nor WS-12 (Cold^+^ AITC^-^ WS-12^-^), and the percentage of CI cells (Cold^-^) from DiI^-^ (left) and DiI^+^ (right) neurons, *** p<0.001, Fisheŕs exact test, DiI^-^ n=477, DiI^+^ n=95 cells. (E) Mean amplitudes of calcium responses to different stimuli, * p<0.05 one-way ANOVA with Sidak’s post hoc test. (F) Graph showing the cumulative number of DiI^-^ (grey) and DiI^+^ (red) CS cells over temperature decrease. (G) Number of neurons responding at the indicated temperature ranges, * p<0.05 Fisheŕs exact test.

Consistent with our previous findings in unlabeled neurons (see above), the number of CS neurons was strongly reduced in Trpa1 KO mice. After DiI application to the lungs, only 1 out of 17 DiI^+^ neurons (5.9%) responded to cold and this reduction was observed in DiI^-^ neurons as well (16 out of 301, 5.3%).

To exclude the possibility that the increase in CS neurons in the lungs or airways was caused by the injection or damage produced by the dye itself, we performed the same analysis after injection of DiI into the stomach wall. These injections marked an average of 2.6 % (16/604) in cultures of the nodose-jugular ganglion complex (not shown). As shown in figure 8A, CS neurons innervating the stomach wall could also be identified in nodose cultures. However, the fraction of CS neurons was small (17.8%), a significantly lower percentage when compared to the lungs (35.8%) (p = 0.0317, Fisher exact test). Unlike CS neurons innervating the lungs, in the stomach, the fraction of CS neurons (figure 8B) and their response amplitude to cold and AITC was similar to that found in unlabeled neurons (figure 8E). In agreement with the characteristics of lung-innervating neurons, only a small fraction responded to WS-12 (figure 8B,D). Noteworthy, the same percentage (50%) of CS neurons was responsive or not to AITC. This result, together with the low number of WS-12 responses, suggests the presence of a significant population of CS neurons innervating the stomach that is independent of TRPA1 and TRPM8.

**Figure 8.**
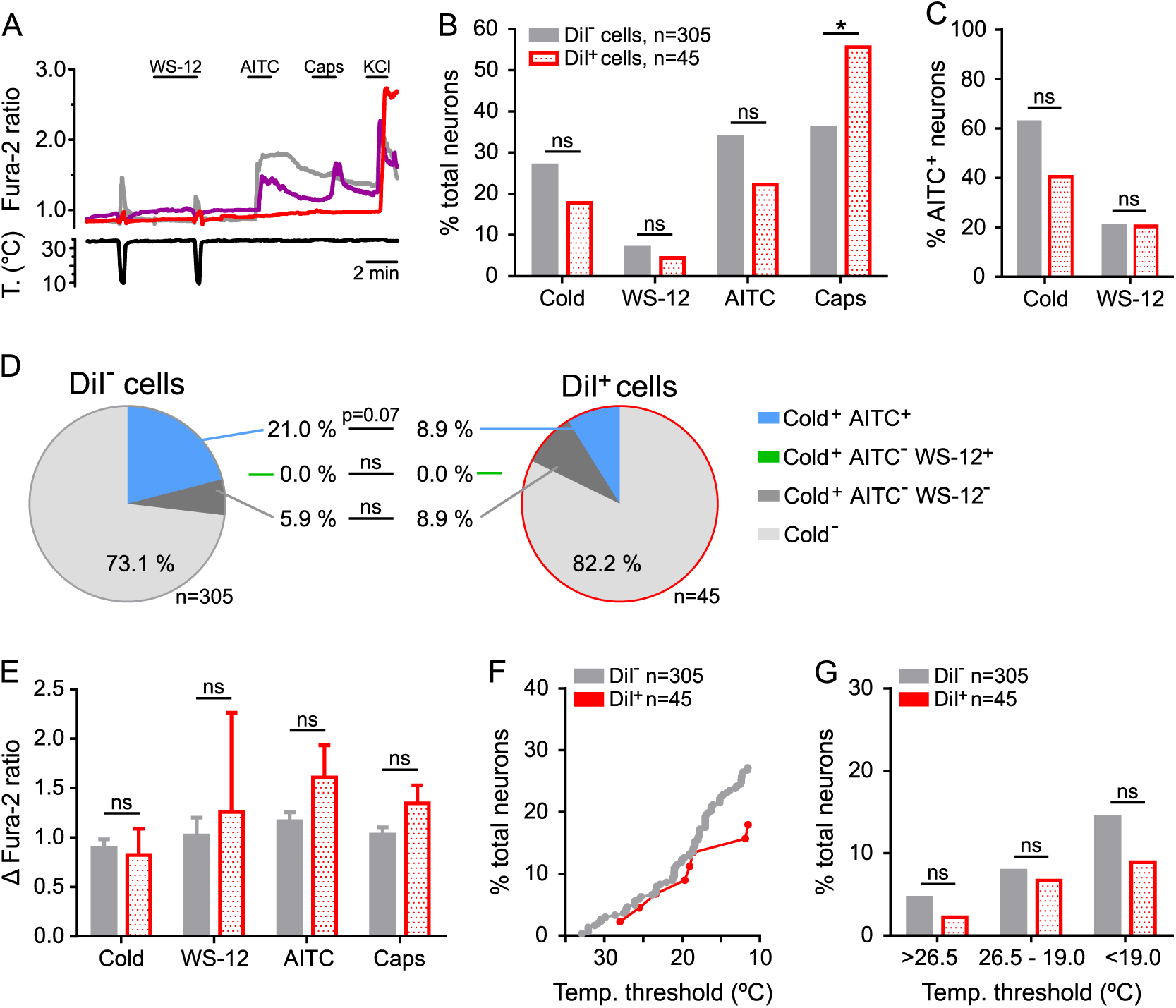
Phenotypic characterization of stomach-innervating CS vagal neurons. (A) Exemplary traces of two DiI^+^ (red and purple) and one DiI^-^ (grey) neurons responding to different stimuli combinations: cold, 500 nM WS-12, 100 µM AITC, 100 nM Capsaicin, 50 mM KCl. (B) Percentage of total number of DiI^+^ and DiI^-^ neurons responding to each stimulus, * p<0.05, Fisheŕs exact test. (C) Number of cells responding to cold or WS-12 out of all AITC-sensitive neurons, * p<0.05, Fisheŕs exact test, DiI^-^ n=134, DiI^+^ n=39 cells. (D) pie charts showing the percentage of total cells that respond to cold and AITC (Cold^+^ AITC^+^), WS-12 but not AITC (Cold^+^ AITC^-^ WS-12^+^), or neither AITC nor WS-12 (Cold^+^ AITC^-^ WS-12^-^), and the percentage of CI cells (Cold-) from DiI^-^ (left) and DiI^+^ (right) neurons, Fisheŕs exact test, *** p<0.001, DiI^-^ n=305, DiI^+^ n=45 cells. (E) Mean amplitudes of calcium responses to different stimuli, one-way ANOVA with Sidak’s post hoc test. (F) Graph showing the cumulative number of DiI^-^ (grey) and DiI^+^ (red) CS cells over temperature decrease. (G) Number of neurons responding at the indicated temperature ranges, Fisheŕs exact test.

## Discussion

### Distinct cold sensing mechanisms in mouse VG and TG

Brain function and behavior are shaped by the constant evaluation of external and interoceptive information (Critchley & Harrison, 2013; Hamed et al., 2024). Temperature sensing by cutaneous and visceral afferents participate in these regulatory loops but their molecular mechanisms and circuit pathways are still poorly understood. We performed a side-by-side comparison of the involvement of TRPA1 and TRPM8 as cold transducers in mouse vagal and trigeminal ganglia and note major differences between them.

Applying a cold ramp from ∼33 to ∼11 °C, (∼30 seconds to reach the lowest temperature), we found a variety of cold-sensitive (CS) neurons in both ganglia, with different temperature thresholds, amplitudes and response profiles. Our data showed that 22 % of VG neurons responded to cold. Fajardo et al. found a percentage of 17 % in a smaller sample of mouse VG neurons, using a similar cold ramp. Interestingly, they also found a much higher percentage of CS neurons in rats (49 %), suggesting a difference between the two rodent species (Fajardo et al., 2008). In contrast to VG, there were only 15 % of CS neurons in TG. Previous studies reported a wide range in the number of trigeminal CS neurons. For example, Viana et al. found 9 % of CS neurons when lowering the bath temperature to ∼15 °C (Viana et al., 2002). Similarly, McKemy et al reported 13.5 % CS neurons for brief cold pulses to 7 °C (McKemy et al., 2002). In contrast, Karashima found 23 % of CS neurons in TG, using a cold ramp reaching ∼10 °C. However, the dynamics of the cold ramp were very different in this study, being slower and maintained for much longer (5 min) (Karashima et al., 2009), which might explain the increased number of responders. These differences in experimental protocols and outcomes highlight the importance of the side-by-side comparison we performed, to understand the role of different channels in specific physiological context.

To compare TRP channel expression and their involvement in mediating cold responses in VG and TG neurons, pharmacological (TRP agonists and antagonists), biochemical (qPCR) and genetic (i.e. neurons from knock out mice) approaches were used. In VG, the great majority of CS neurons also responded to the TRPA1 agonist AITC, were blocked by the TRPA1 antagonists HC-030031 and A967079, and were nearly absent in cultures from Trpa1 KO mice. A previous report showed that AITC can also activate TRPM8 channels and this could potentially affect our classification scheme. However, TRPM8 activation by AITC requires millimolar concentrations, one or two orders of magnitude higher than the concentration we used (Janssens et al., 2016).

Altogether, the data clearly indicate that TRPA1 is the main cold transducer in mouse vagal ganglion neurons, covering the whole temperature range. This agrees with previous results (Fajardo et al., 2008). In alignment with the very low expression of TRPM8 in VG shown in this study as well as by others (Kupari et al., 2019; Meerschaert et al., 2020; Nassenstein et al., 2008; Yu et al., 2015), we further showed that TRPM8 plays a minor role in the mechanism of cold transduction in mouse VG, a role that is probably restricted to the jugular part of the vagal ganglion complex.

In trigeminal CS neurons the contribution of TRPA1 and TRPM8 were quite different, with only 23 % of CS neurons responding to AITC (TRPA1-like), but 59 % to menthol and not AITC (TRPM8-like). Importantly, the data clearly showed that in TG all the cold responders with large Fura-2 transients and low temperature thresholds were of the TRPM8-like profile (menthol^+^/AITC^-^). Furthermore, in neurons from Trpm8^-/-^ mice, the responses with large amplitudes and low thresholds were missing in the AITC^-^ CS (TRPM8-like) group. The data further showed that the highly selective TRPM8 antagonist RQ00203078 reduced the fraction of cold responders by more than half in TG. Additionally, cultures from Trpm8^-/-^ mice showed a very similar reduction of CS neurons, and this reduction was present in the AITC^-^ CS neurons, which contain the TRPM8^+^ CS neurons. This agrees with previous *in vitro* studies showing that TRPM8, but not TRPA1, mediates low-threshold cold responses in TG (Bautista et al., 2007; Madrid et al., 2009; McKemy et al., 2002). In contrast to the TRPM8-mediated cold responses, the AITC-sensitive CS neurons (TRPA1-like) showed much smaller amplitudes and higher temperature thresholds; notably only 4/17 neurons responded to a temperature above 20 °C. In line with this finding, there was a conspicuous absence of cold responses below 18 °C in cultures from Trpa1^-/-^ mice, showing the importance of TRPA1 in sensing noxious but not mild cold temperatures in TG. This was very different in VG, where AITC^+^ CS neurons covered the whole range of amplitudes and temperature thresholds.

Therefore, the data clearly show that TRPM8 mediates the majority of cold responses in TG and all responses to mild temperature drops, whereas TRPA1 plays some role in mediating additional high threshold cold responses. The involvement of TRPM8 as a cold transducer in TG neurons has long been established, especially for low-threshold (>20 °C) thermoreceptors (Bautista et al., 2007; Leijon et al., 2019; Madrid et al., 2009; McKemy et al., 2002; Thut et al., 2003). *In vivo* electrophysiological recordings from trigeminal nucleus caudalis in mice identified neurons activated by mild cooling of the oral cavity and others strongly activated by noxious cold (<10 °C) (Li et al., 2024). Notably, in TRPM8 KO mice robust responses to cold temperatures persisted but, the great majority were classified as high threshold and menthol insensitive.

Evidence for the contribution of TRPA1 channels in cold sensing has been inconsistent and even sensitive to subtle changes in the testing paradigm employed (Winter et al., 2017). Some studies found that TRPA1 did not play a role in trigeminal cold sensing, even when lowering the temperatures to below 10 °C (Bautista et al., 2006; Jordt et al., 2004; Madrid et al., 2009; Yarmolinsky et al., 2016). However, most of these studies (Bautista et al., 2006; Jordt et al., 2004; Madrid et al., 2009) used TG neurons isolated from newborn mice, where TRPA1 expression is lower than in neurons from adult mice (Hjerling-Leffler et al., 2007; Memon et al., 2017). In agreement with our new results, it has been shown before that a percentage of rat TG CS neurons are isolectin B4 (IB4) positive (Thut et al., 2003), which suggests TRPA1 expression (Barabas et al., 2012). Notably, Karashima et al. found that the majority of mouse CS TG neurons were AITC^+^, and that this subpopulation was absent in neurons from Trpa1^-/-^ mice (Karashima et al., 2009). These differences might be explained by the much slower and longer cold ramp (maintained for 5 minutes), compared to the brief ramps used here. Sustained and intense cold stimuli might activate more high-threshold, TRPA1-like CS neurons. In fact, it has been shown recently that a prolonged cooling stimulus (5 min at 1 °C) to the paw of anesthetized mice unmasked a population of late-responding, Nav1.8-positive, CS DRG neurons that only started to respond 100-250 seconds after the start of the cold ramp (Luiz et al., 2019). This population of CS neurons might have remained undetected in many studies using short stimuli, including the data presented here.

Adults VG and TG had a significant proportion of CS neurons that did not respond to either menthol or AITC (29 % in VG and 19 % in TG), suggesting the presence of a hitherto unknown TRPA1-/TRPM8-independent transducing mechanism. This population was also confirmed in neurons from double KO mice. In VG, these TRPA1-/TRPM8-independent CS neurons generally have small amplitudes, as previously reported for Trpa1 KO mice (Fajardo et al., 2008). Furthermore, they respond to a wide range of temperature thresholds. In TG, CS neurons with an unknown transducer mechanism showed generally small amplitudes and higher temperature thresholds, similar to TRPA1-mediated CS neurons.

Few studies have performed a functional characterization of cold responses in mice lacking both TRPM8 and TRPA1. Previous studies in DRG and TG neurons have shown, using pharmacological tools, that there is a proportion of neurons that mediate cold responses via an unknown, TRPA1-/TRPM8-independent mechanism (Babes et al., 2004; Leijon et al., 2019; Memon et al., 2017; Munns et al., 2007). The molecular identity of this additional cold sensor remains to be characterized. TRPC5, another cool-activated TRP channel, has been proposed to mediate some of the cold responses in DRG neurons, especially at mild cold temperatures (>25 °C) (Zimmermann et al., 2011). TRPC5 is expressed in mouse TG (Gomis et al., 2008) as well as in rat nodose ganglia (Zhao et al., 2010). However, single cell RNA sequencing showed no TRPC5 expression in mouse VG (Kupari et al., 2019). Recently, TRPC5 has been shown to be highly expressed in odontoblasts and to be an important contributor to dental cold sensitivity (Bernal et al., 2021). However, our data do not support a significant role of TRPC5 in cold sensitivity in isolated TG and VG because most of the TRPA1-/TRPM8-independent neurons are high threshold CS neurons, being activated by temperatures lower than 25 °C. Recently, two studies showed that glutamate receptor GluK2 is important for noxious cold (10 °C) but not cool (22 °C) sensing in mouse DRG neurons (Cai et al., 2024; Gong et al., 2019). RNA sequencing of TG shows expression of the GluK2 gene Grik2 (Nguyen et al., 2017), making it a possible candidate as a noxious cold transducer in the CS neuronal population described here.

In summary, in this side-by-side comparison of VG and TG cold-sensitive neurons, we confirmed that the majority of vagal CS neurons are mediated via TRPA1, covering the whole range of temperature thresholds. In contrast, a large proportion of trigeminal CS neurons are mediated via TRPM8, mainly low threshold cold responses. TRPA1 plays a minor role, mainly in high threshold CS neurons. These functional differences in cold sensitivity find a molecular correlate in recent transcriptomic studies indicating a strong segregation in the molecular profile of somatic (DRG, TG) and vagal neurons (Kupari et al., 2019; Meerschaert et al., 2020; Nguyen et al., 2017; Zeisel et al., 2018), consistent with their different developmental origin (Baker, 2005). Our reanalysis of published single-cell transcriptomic data for mouse TG and VG found that TG had fewer TRPA1^+^ neurons than VG (13.3% *vs* 19.8%), and their expression level, calculated as reads per million (RPM) for mean Trpa1 expression normalized to RMP values for actin beta (Actb), was also higher in VG (0.558 *vs* 0.217).

### Different sensitivity to cold in TRPA1-expressing VG and TG neurons

Our calcium imaging data show that the number of TRPA1-expressing neurons (AITC^+^) is similar between VG and TG. However, the percentage of AITC^+^ neurons that also respond to cold is much higher in VG (63 %) than in TG (11 %) neurons, showing that visceral TRPA1-expressing neurons are more sensitive to cold than their trigeminal counterparts. Like our results in TG, previous studies found that CS DRG and TG neurons make up between 8 and 22 % of AITC^+^ neurons (Madrid et al., 2009; Memon et al., 2017; Munns et al., 2007). Small differences in the exact percentages can be explained by experimental variables, including different temperature ramps, or the age of mice, which influence the TRPA1 expression level (Hjerling-Leffler et al., 2007; Memon et al., 2017). So, what are the molecular differences between visceral and somatosensory neurons that make visceral TRPA1-expressing neurons more sensitive to cold? One simple explanation is the expression of higher levels of TRPA1. First, the amplitude of calcium transients in AITC^+^ neurons was larger in VG than in TG neurons. Previously, Memon et al. also showed that CS AITC^+^ DRG neurons express more TRPA1 compared to cold-insensitive (CI) AITC^+^ neurons (Memon et al., 2017). In alignment with this finding, we found that the calcium transients to AITC were larger in CS neurons than those in CI neurons. This was the case in both ganglia. In addition, we found that the Trpa1 RNA expression level was higher in VG compared to TG neurons. TRPA1 is a calcium permeable channel and previous studies described a complex positive feedback loop of permeant calcium on TRPA1 gating (Doerner et al., 2007). Elevated cytosolic calcium has a bimodal effect, it potentiates TRPA1 activity at low concentrations and inactivates the channel at higher Ca^2+^ concentrations (Hasan et al., 2017; Y. Y. Wang et al., 2008; Zurborg et al., 2007). Thus, higher TRPA1 expression could lead to a transient potentiation of TRPA1 during cooling. Therefore, a higher density of TRPA1 channels per TRPA1-expressing neuron in VG could explain why these neurons are more sensitive to cold. Other potential explanations for the differences between VG and TG include differences in the glycosylation status (Egan et al., 2016), or their localization in restricted membrane domains (Startek et al., 2019). Both have been shown to influence TRPA1 agonist sensitivity and surface expression.

An additional possibility to explain the different sensitivities in TRPA1^+^ neurons between VG and TG is the negative regulation of the neuron’ excitability by potassium channels. The responsiveness of trigeminal TRPM8-expressing neurons to a cold stimulus is not only governed by the expression level of TRPM8 channels but also by the expression levels of K_v_1.1/1.2 channels that act as a brake on their excitability (Madrid et al., 2009). More recently, a similar suggestion has been made for the regulation of TRPA1^+^ neurons by K_v_1.2 channels in DRG neurons (Memon et al., 2017). Our qPCR data show that K_v_1.1 is expressed at similar levels in both VG and TG ganglia. Single cell RNA sequencing from vagal neurons revealed that TRPA1-expressing clusters co-express *Kcna2*, the gene for Kv1.2, but not *Kcna1* (Kv1.1) (Kupari et al., 2019), possibly suggesting a regulatory role of Kv1.2 in TRPA1-expressing neurons. Using the potassium channel blocker 4-AP and calcium imaging on cultured vagal neurons, we further showed that a potassium channel block not only activates silent CS neurons but also sensitizes original CS neurons to cold temperatures. This could be due to an increased neuronal excitability after removal of its brake. However, these two phenomena were still present in neurons from Trpa1^-/-^ neurons, suggesting the mechanism to be independent of TRPA1 expression. Furthermore, other potassium channels have been shown to be expressed on mouse and rat vagal sensory neurons, including TRESK, TREK1, TREK2, TRAAK, TASK-2, TASK-3, TWIK, Kv1.4, and Kv4 (Cadaveira-Mosquera et al., 2012; Matsumoto et al., 2011; Park et al., 2018; Zhao et al., 2010). Most of these channels have also been shown to influence cold sensitivity (Castellanos et al., 2020; Morenilla-Palao et al., 2014; Noel et al., 2009). This raises the possibility that any of these potassium channels could dampen the excitability of TRPA1-expressing neurons. However, functional studies investigating the different expression levels in cold-sensitive and cold-insensitive neurons are required before any further conclusions can be drawn.

### Low and restricted TRPM8 expression in VG

Using different approaches (qPCR, immunohistochemistry, pharmacological interrogation and calcium imaging in transgenic mice), we found a lower expression and reduced functionality of TRPM8 in VG compared to TG. First, our calcium imaging data showed that only 0.7 % of all vagal neurons responded to menthol but not AITC, which was used as an indication of menthol activating TRPM8 instead of TRPA1. The lack of reduction in the number of CS neurons in cultures from Trpm8^-/-^ mice further indicates low TRPM8 expression in VG. In a recent study, using a transgenic mouse line that expresses the fluorescent protein EYFP in TRPM8^+^ neurons (Morenilla-Palao et al., 2014), we found that ∼18% of TG neurons express EYFP (Alcalde et al., 2018). Here, using the same mouse line, the number of labelled neurons in VG was only 4.5 %. The VG complex originates from two embryologically distinct neuronal populations: neural crest-derived jugular ganglion (JG) as well as placode-derived nodose ganglion (NG) neurons (Baker, 2005). We found that the majority of the few TRPM8-expressing vagal neurons are located in the JG, the somatic part of the ganglion complex. This corresponds with recently published single cell RNA sequencing results from mouse VG neurons. The authors used the neural crest marker Prdm12 to distinguish JG from NG neurons. TRPM8 transcripts were found in only 1 out of 6 JG clusters, which accounts for ∼12 % of all JG neurons (Kupari et al., 2019). Similarly, using *in situ* hybridization experiments, it has been shown previously that TRPM8 expression is limited to the JG portion of the VG in rats (Hondoh et al., 2010). Functional and molecular studies in guinea pig vagal complex fully support the restricted expression of TRPM8 to the JG (Yu et al., 2015). Notably, in larger mammals, the JG is ∼20 % the size of the NG (Mazzone & Undem, 2016) and only ∼20 % of mouse VG neurons express the JG marker Prdm12, whereas ∼80 % express the NG markers Phox2b and P2rx2 (Kupari et al., 2019; Prescott et al., 2020). Similarly, our immunohistochemical data showed that 74 % of all vagal neurons are part of the NG and only 26 % make up the JG. Overall, our findings support and expand previous data indicating low and restricted TRPM8 expression in VG, using a variety of techniques (qPCR, RNA sequencing and *in situ* hybridization) (Hondoh et al., 2010; Kupari et al., 2019; Nassenstein et al., 2008). Data presented in this study, including functional experiments, confirm these results.

The homeoprotein Phox2b is as master regulator of visceral circuits, commanding a somatic-to-visceral switch in cranial sensory nuclei (D’Autreaux et al., 2011). It would be interesting to investigate the functional phenotype of vagal neurons deficient for Phox2b. We predict that they will show cold responses similar to those observed in TG.

### Enhanced cold sensitivity in airway-innervating vagal neurons

Lastly, we investigated the cold sensitivity of vagal sensory neurons that innervate different visceral territories. We found that CS neurons are overrepresented in vagal neurons innervating the lungs and lower airways, and that the great majority of these CS neurons respond to the TRPA1 agonist AITC. This stands in contrast to an *ex vivo* study using a rat lung preparation which showed that CS bronchopulmonary vagal neurons respond to menthol, but not to cinnamaldehyde, suggesting the involvement of TRPM8 (Zhou et al., 2011). We already mentioned major differences in cold sensitivity of vagal neurons between rats and mice (Fajardo et al., 2008). Our data in mice do not support a major involvement of TRPM8 on vagal airway-specific neurons as only 1 % of them showed a TRPM8-like response profile (WS-12^+^ and AITC^-^). Furthermore, a recent study using single cell RNA sequencing from mouse airway-innervating vagal neurons did not detect TRPM8 transcripts in any of the 72 studied cells (Mazzone et al., 2020). On the other hand, TRPA1 transcripts were found in most neurons. In agreement with the notion that trigeminal and vagal territories show distinct innervation patterns, a recent study described the absence of TRPM8 fibers innervating the mouse trachea, in contrast with their innervation of the nasal mucosa (Jiang et al., 2024).

To fulfill their different functions, interoceptive signals from different organs may have organ-specific features (Wang & Chang, 2024). The vagus nerve innervates various internal organs but the airways and the lungs stand out because they are exposed to external temperature fluctuations. The final air temperature that reaches the lower airways depends on the ambient temperature as well as on the breathing rate (D’Amato et al., 2018). Under extreme conditions, such as athletes exercising at subfreezing temperatures, the drop in inhaled airway temperature can be substantial (Cruz & Togias, 2008). In our data, a large fraction of CS airway-innervating vagal neurons responded to low temperatures (<19 °C) *in vitro*. One may question their physiological relevance as cold sensors. However, cold air is also dry, leading to water loss and hyperosmolarity of the epithelial surface. Besides cold temperature activation, TRPA1 is a polymodal sensor, responding to a wide range of chemical irritants, endogenous agonists (reviewed by Talavera et al., 2020; Viana, 2016) and mechanical forces (Moparthi & Zygmunt, 2020). Gating of TRPA1 by agonists, including temperature, is very complex and sensitive to the lipid environment and redox state (Moparthi et al., 2022; Zimova et al., 2020). Thus, these vagal endings may function as polymodal receptors, engaged in the detection of irritant chemical stimuli in addition to cold temperatures. Infectious agents and other inhaled irritants like cigarette smoke, chlorine, tear gas, or ozone, have been shown to activate TRPA1 (Andrè et al., 2008; Bessac et al., 2009; Kichko et al., 2015; Meseguer et al., 2014; Taylor-Clark & Undem, 2010). Also, nitrative stress and reactive oxygen species have been shown to activate TRPA1 on vagal sensory neurons as well as being implicated in airway hypersensitivity (Bessac et al., 2008; Ruan et al., 2014; Taylor-Clark et al., 2009). In addition, many metabolites and inflammatory products, including bradykinin, LPS, oxidized phospholipids, methylglyoxal, acrolein and ATP, can sensitize TRPA1 activity (Andersson et al., 2013; Bandell et al., 2004; Bautista et al., 2006; Liu et al., 2016; Meseguer et al., 2014; Trevisani et al., 2007; S. Wang et al., 2008). Therefore, the release of a myriad metabolites during lung infection or inflammation may enhance the sensitivity of vagal endings to cold temperatures, exposing a major role of TRPA1 in pathological states (Balestrini et al., 2021; Chen et al., 2011; del Camino et al., 2010; Nassini et al., 2014).

## Materials and Methods

### Animals

Studies were performed on young adult (6-12 weeks old) male and female mice. Mice were bred at the Universidad Miguel Hernández Animal Research Facilities (ES-119-002001) and kept behind a specific pathogen free barrier under 12/12 h light dark cycle with food and water *ad libitum*. Wild type animals were of the C56Bl6/J strain (Envigo/Harlan). All experimental procedures were performed according to the Spanish Royal Decree 1201/2005 and the European Community Council directive 2010/63/EU, regulating the use of animals in research. The following transgenic mouse lines were used for calcium imaging experiments: The TRPA1 KO line was generated based on mice kindly provided by Dr. D. Corey (Harvard Medical School, Boston,MA) (The Jackson Laboratory stock no. 006401) (Kwan et al., 2006). The TRPM8 KO mice were based on a transgenic knockin line, Trpm8^EGFPf^ (Dhaka et al., 2007). To generate the Trpa1::Trpm8 double KO line, Trpm8 KO mice with enhanced EGFPf expression (Dhaka et al., 2008) were crossed with the above mentioned Trpa1 KO line. Transgenic mice were backcrossed into the C56Bl6/J background for at least 5 generations. Mice with a wild type genotype from all three transgenic lines were pooled and used as controls for the experiments with KO mice. Trpm8^BAC-EYFP^ mice were used for histology experiments to quantify TRPM8^+^ neurons in VG (Morenilla-Palao et al., 2014). The genotype of transgenic mice was established by PCR.

### Retrograde labelling of lung afferents

Airway-innervating vagal sensory neurons were retrogradely labelled with 1 % 1,1’-dioctadecyl-3,3,3,’3-tetramethylindocarbocyanine methanesulfonate (DiI) (DiIC18(3), Thermo Fisher Scientific; 10 % stock in DMSO, diluted in physiological saline) or 5 mg/ml WGA-Alexa594 (Thermo Fisher Scientific; diluted in physiological saline), following the method presented by Vandivort (Vandivort et al., 2016). Mice were anaesthetized with isoflurane for 2-3 minutes and then orotracheally intubated with a home-made catheter consisting of a 3-4 cm long polyethylene 20G piece of tubing attached to a 21 G needle. DiI or WGA-Alexa594 was introduced in a volume of 50 µl followed by ∼300 µl of air to make sure the dye spreads throughout the distal airways and into the alveoli. Mice were returned to their cages, recovered within a few minutes from anesthesia, and sacrificed 3-7 days after intubations for tissue extraction.

### Retrograde labelling of stomach afferents

Stomach-innervating vagal sensory neurons were retrogradely labelled with 0.5 or 1 % DiI prepared from a 5 or 10 % stock in DMSO and diluted in physiological saline. Mice were anaesthetized with isoflurane for 2-3 minutes and injected with buprenorphine (0.1 mg/kg s.c). Thereafter, the stomach was surgically exposed through a laparotomy. Each mouse received 10 small injections (∼0.25 µl) of DiI in the stomach wall (5 on the dorsal side and 5 on the ventral side) with a 5 µl Hamilton syringe. The wound was sutured with absorbable n° 6 suture internally and Michel clips externally. Mice were returned to their cages, and administered paracetamol orally for 2-4 days post-surgery, before being sacrificed 3-7 days after surgery for tissue harvest.

### Isolation and culture of VG and TG neurons

Mice were killed by CO_2_ and the heart was cut to confirm death. By gaining access via the ventral aspect of the neck and following the vagus nerve, the vagal ganglia (VG) were located and carefully cut at the rostral and then the caudal end. When possible, trigeminal ganglia (TG) were removed from the same mice immediately after VG dissection to reduce the number of animals used. The skull was opened, the brain was removed, and TG were carefully cut out. Excised ganglia were immediately placed in ice cold HBSS, any remaining nerve fibers were trimmed, and TG were cut into 2-3 pieces. In two mice, used to retrogradely label lung afferents, the DRG ganglia from different spinal levels were also removed.

Ganglia were dissociated enzymatically (collagenase type XI (1800 U/ml, Sigma) and dispase II (5 U/ml, Gibco) for around 45 min at 37 °C in 5 % CO_2_) and mechanically (10-15 times pipetting with 1 ml plastic pipette tip), filtered through a 70 µm nylon colander (Falcon) with 3-4 ml of Ca^2+^- and Mg^2+^-free HBSS medium containing 10 % fetal bovine serum (FBS, Gibco), 1 % MEM-vit (Gibco) and 1 % penicillin/streptomycin (Gibco), and centrifuged at 300 g for 10 minutes. For TG and DRG, the pellet was resuspended in Minimum Essential Medium (MEM, Gibco) supplemented with 10 % FBS, 1% MEM-vit and 1% penicillin/streptomycin. Since there was no visible pellet for VG, as much medium as possible was aspirated without disturbing the cells at the bottom of the Falcon tube and cells were resuspended in the leftover medium. TG and VG cells were plated on poly-L-lysine (0.01 %, Sigma) coated glass coverslips (Ø 12 mm, thickness no 1, Marienfeld Superior) and left for ∼1 hour to adhere to the coverslips before being flooded with ∼300 µl MEM supplemented with 10 % FBS, 1% MEM-vit and 1% penicillin/streptomycin. No additional growth factors were added. Calcium-imaging recordings were performed after 4-24 hours in culture.

### Calcium microfluorimetry

Ratiometric calcium imaging experiments were conducted with the fluorescent indicator Fura-2 (Thermo Fisher Scientific or Biotium). Neurons were incubated with 5 μM Fura-2 AM and 0.04% Pluronic F-127 (Thermo Fisher Scientific) for 45-60 min at 37 °C in standard extracellular solution. Fluorescence measurements were obtained on a Nikon Eclipse TE2000-U inverted microscope fitted with an Andor 888 camera (Oxford Instruments) and a Nikon 20x S Fluor (NA 0.75) objective. Fura-2 was excited at 340 and 380nm with a rapid switching Polychrome V monochromator (TILL Photonics) and mean fluorescence intensity ratios (F340/F380) were displayed with Life Acquisition software (FEI Munich GmbH) every second. The standard extracellular solution contained (in mM): 140 NaCl, 3 KCl, 2.4 CaCl_2_, 1.3 MgCl_2_, 10 HEPES, and 10 glucose, and was adjusted to a pH of 7.4 with NaOH. Calcium imaging experiments were performed at a basal temperature of 33 ± 1 °C. Before the start of the experiment, an image of the microscopic field was obtained with transmitted light. For experiments using the retrograde dye DiI, another image was taken under 550 nm excitation wavelength in order to identify labeled neurons. Regions of interest (ROIs) were manually defined around single neurons. Custom-written Matlab scripts were used to quantify responses to cold and chemical agonists. Responses were scored as positive if the increase in fluorescence (ΔFura2 ratio) was > 0.05. Neurons were considered healthy if they responded to 50 mM KCl applied at the end of each protocol. Cells that failed to respond to KCl but showed clear responses to AITC or capsaicin were also considered viable neurons. Only cells with a diameter > 5 µm were analyzed.

### Temperature stimulation

Coverslips with cultured cells were placed in a microchamber and continuously perfused with solutions at 33 ± 1°C. The temperature was adjusted with a Peltier device (CoolSolutions, Cork, Ireland) placed at the inlet of the chamber and controlled by a feedback device (Reid et al., 2001). Cold sensitivity was investigated with a temperature drop to 11.5 ± 1 °C (∼ -1.5 °C/s). The bath temperature was monitored with an IT-18 T-thermocouple connected to a Physitemp BAT-12 thermometer (Physitemp Instruments) and digitized with an Axon Digidata 1440A converter and Clampex 10 software (Molecular Devices). Temperature and cellular calcium signals were recorded simultaneously. The temperature threshold for each neuron was measured manually, estimated from the time point when calcium level starts to increase.

### Calcium imaging protocols

The data shown in figure 1 were acquired in three different sets of experiments with slightly different protocols. All ganglia were extracted and cultured from wild type mice following the same protocol as described above. A cold ramp was applied as the first stimulus in all three protocols and the data in figure 1 corresponds to responses to this first stimulus only. The number of neurons responding to cold and their amplitudes were similar between the different sets for VG and TG (Table 1). For the last set (Set 3), ganglia were extracted from male mice that had been used as a control group in a cohort of experiments not included in this manuscript. In this case, mice were injected intraperitoneally three times with 5 % glucose in distilled water on alternate days, and the ganglia were extracted 4-5 days afterwards. The data shown in figures 2 and 6 are also from this set of experiments.

**Table 1.** Summary of data pooled from three different sets of experiments. The percentage of responses to a cold ramp and their mean response amplitudes were similar in the different experiments for VG and TG, one-way ANOVA with Tukeýs post hoc test. Number of neurons for each set of experiments (number of coverslips are shown in brackets): VG Set 1 n=531 (18), VG Set 2 n=388 (17), VG Set 3 n=402 (11), TG Set 1 n=213 (14), TG Set 2 n=243 (17), TG Set 3 n=597 (17).

| | Set1<br>Mean ± SEM | Set 2<br>Mean ± SEM | Set 3<br>Mean ± SEM |
| --- | --- | --- | --- |
| **VG CS neurons (%)** | 19,33 ± 1,96 | 21,98 ± 2,75 | 23,60 ± 3,56 |
| **TG CS neurons (%)** | 15,11 ± 1,85 | 16,88 ± 2,55 | 13,01 ± 1,45 |
| **VG cold response (DF)** | 0,60 ± 0,08 | 0,65 ± 0,08 | 0,39 ± 0,05 |
| **TG cold response (DF)** | 0,65 ± 0,14 | 0,83 ± 0,13 | 0,53 ± 0,06 |

For experiments with TRPA1 (HC030031 (HC) or A967079 (A96)) or TRPM8 (RQ00203078 (RQ)) antagonists (Set 1), two cold ramps were applied: the first one either in control conditions (extracellular solution) or in the presence of the antagonist after a 100 sec pre-incubation; the second one after a washout time of 3 min (ctrl, HC) or 6-7 min (A96, RQ). The second cold ramp was used to determine which neurons were cold-sensitive. Some additional experiments were performed with the TRPM8 antagonist RQ00203078, adding a cold ramp in control conditions in the beginning of the protocol. This first cold ramp was used to determine which neurons were cold-sensitive in these experiments. For experiments with the potassium channel blocker 4-Aminopyridine (4-AP) (Set 2), two cold ramps were applied: the first one in control conditions and the second one in the presence of 4-AP after a 100 sec pre-incubation. In a fraction of these experiments, WS-12 and AITC was applied after the two cold ramps to obtain the data shown in figure 5.

### Magnetic-activated cell sorting (MACS)

VG and TG were dissected and cleaned from fibers as described above. They were incubated in collagenase type XI (2700 U/ml, Sigma) and dispase II (5 U/ml, Gibco) at 37 °C in 5 % CO_2_ for 20 minutes followed by brief mechanical dissociation. Cells were passed through a 40 µm filter (Falcon), washed with 4-5 ml ice cold PBS supplemented with 0.5 % BSA (Tocris), and collected by centrifugation for 10 min (300 g). Myelin was removed with the Myelin Removal Beads II kit (Miltenyi Biotec, 130-096-731) followed by the separation of neurons from non-neuronal cells (astrocytes, oligodendrocytes, microglia, endothelial cells, and fibroblasts) using the Neuron Isolation Kit (Miltenyi Biotec, 130-115-389) following the kit’ instructions. A pure population of neurons was obtained, resuspended in Trizol (Ambion, Life Technologies, 15596026) and stored at -80 °C.

### Quantitative PCR

RNA was extracted from MACS sorted neurons (see above) using the RNeasy Micro Kit (Qiagen, 74004) following the manufactureŕs instructions. Extracted RNA was converted into complementary DNA (cDNA) and amplified by RT-PCR using SuperScript III Reverse Transcriptase (Thermo Fisher Scientific) and a random primer mix (Thermo Fisher Scientific). Quantitative PCR (qPCR) was performed in triplicates using 1:5 diluted cDNA, the PyroTaq EvaGreen qPCR Master mix (Cultek Molecular Bioline) and the following primers (Sigma, 5′-3′): 18S F: GCAAGACGGACCAGAGCGAAAG, R: ATCGCCAGTCGGCATCGTTTATG; β-actin F: TTCTTGGGTATGGAATCCTGTGG, R: GAGGTCTTTACGGATGTCAACG; TRPM8 F: AAGCTCGTTGTACTCTGGGC, R: AAAACGTCGATCAGCATCCG; TRPA1 F: GCTTCACAGAGCCTCGTTATTTG, R: TGCTGTTGATGTCTGCTCCC; TRPV1 F: CCCCAAGGCTCTATGATCGC, R: TTCCCGTCTCTGGGTCTTTG; Kv1.1 F: CCATGACCACTGTGGGATACG, R: GCCTCCAACTGTCACAGGG. Threshold cycle (Ct) values were determined by setting the threshold to 0.2 and the Ct values of each gene of interest were subtracted from the Ct values of the two housekeeping genes 18S and beta actin (ΔCt). To account for the logarithmic scale during qPCR, results are presented as 2^(-ΔCt)^ values for each gene for each mouse.

### Cytochemistry of cultured cells

VG, TG and DRG (separated into lumbar, thoracic, and cervical ganglia) were dissected from mice 3-7 days after DiI labeling, cultured and placed on glass coverslips as described above. The following day, cells were fixed with 4 % paraformaldehyde (PFA, Scharlau) for 10 min at room temperature, washed with PBS and mounted on microscope slides (Normax) using Vectashield mounting medium (Vector Laboratories, Burlingame, USA). Bright field images and images at 555 nm wavelength were acquired using a Leica SPEII upright confocal microscope (LasX software) with a Leica 20x ACS APO (NA 0.6) oil immersion objective. Imaging fields were chosen randomly. The numbers of total neurons and DiI positive neurons were counted manually for VG, TG and DRG.

### Immunohistochemistry of tissue sections

For some of the lung afferent retrograde labelling experiments, VG, TG, and parts of the lung parenchyma and trachea were dissected from C57BL/6J mice 3-7 days after WGA-Alexa 594 intubation. Furthermore, VG were dissected from Trpm8^BAC-EYFP^ mice for the quantification and location of TRPM8-expressing neurons. The endogenous EYFP signal was enhanced with an anti-GFP antibody. An anti-tubulin β III antibody was used in all sections as a marker for neurons. The following antibodies were used (Table 2):

**Table 2:** Primary and secondary antibodies used for immunohistochemistry. Dissected tissues were fixed in 4 % PFA for 2 hours at 4 °C, washed with PBS and cryopreserved with 30 % sucrose at 4 °C overnight. Tissues were mounted with Tissue-Tek (Sakura), cut in 20 µm thick sections using a cryostat (MNT, SLEE Medical) and collected on Superfrost microscope slides (Thermo Fisher Scientific). Slides were dried and washed with 2x PBST (PBS with 0.05% Tween 20) for 10 min on a shaker at room temperature (RT). Blocking buffer was applied for 1 hour at RT, before primary antibody incubation overnight at 4 °C. The next day, tissues were washed with PBST 4x for 15 min on a shaker at RT before being incubated with the secondary antibody for 2 hours at RT. Another 4x 15 min wash with PBST was followed by a 5 min incubation with Hoechst 33342 (Thermo Fisher Scientific), a 10 min wash with PBS and a 5 min wash with distilled water. Tissues were mounted with Fluoromount (Sigma). Images were taken with a Leica SPEII upright confocal microscope (LasX software) with a Leica 20x ACS APO (NA 0.6) oil immersion objective. For detecting WGA-Alexa 594 signals, fluorescence was excited with a 561 nm laser line and collected with an appropriate bandpass filter. Images were analyzed in Fiji (Schindelin et al., 2012). For WGA-Alexa 594 and tubulin β III quantification, positive neurons were counted manually. For the analysis of TRPM8^+^ neurons, the contrast and brightness of the images were increased, and every potentially positive neuron highlighted with an individual ROI. The fluorescence of these ROIs was measured in the original channels for GFP (TRPM8) and normalized to the maximum fluorescent value for each channel (8-bit images, 0-255). Neurons were considered to be GFP^+^ if they had a mean fluorescence value >18 %.

|  | **Name** | **Supplier** | **Reference number** | **Dilution** |
| --- | --- | --- | --- | --- |
| Primary antibody | mouse anti-tubulin $\beta$ III | Biolegend | 801201 | 1:1000 |
|  | chicken anti-GFP | Abcam | 13970 | 1:2000 |
| Secondary antibody | goat anti-mouse Alexa 647 | Invitrogen | A21237 | 1:1000 |
|  | donkey anti-chicken A488 | Jackson IR | 703-545-155 | 1:800 |

### Software and statistics

Data are reported as mean ± standard error of the mean (SEM). When comparing two means, statistical significance (p<0.05) was assessed by Fisheŕs exact test (percentage of responders) or Student’s two-tailed t-test. For multiple comparisons of means, one-way ANOVA with Tukey’s or Sidak’s post-hoc test was performed. Data analysis was performed using Matlab (MathWorks), Excel (Microsoft) and GraphPad Prism version 6.01 for Windows (GraphPad Software, La Jolla California USA). All statistical analysis was performed using GraphPad.

### Chemicals

Menthol (Scharlau), HC-030031 (Tocris) and A967079 (Tocris) were prepared as stocks and stored at 4 °C. AITC (Allyl isocyanate, Sigma), Capsaicin (8-Methyl-N-vanillyl-*trans*-6-nonenamide, Sigma), RQ00203078 (Tocris) and WS-12 ((1*R*\*,2*S*\*)-*N*-(4-Methoxyphenyl)-5-methyl-2-(1-methylethyl)cyclohexanecarboxamide, Alomone) were prepared as stocks and stored at -20 °C. All stocks were diluted in extracellular solution to obtain the final working concentration (see Table 3).

**Table 3.** Agonists and antagonists used in calcium imaging experiments. Square brackets stand for concentration. * A 100 mM stock of capsaicin was prepared in ethanol, and then diluted to a 0.1 mM stock in distilled water (dH2O). ** A 100 mM stock of RQ00203078 was prepared in DMSO, and then diluted to a 0.1 mM stock in dH2O.

| Name | [stock] (mM) | Stock solvent | [final] (μM) | Duration of application (sec) |
| --- | --- | --- | --- | --- |
| Menthol | 300 | DMSO | 100 | ~150 |
| WS-12 | 20 | DMSO | 0.5 | ~150 |
| AITC | 10 | DMSO | 100 | ~150 |
| Capsaicin | 0.1* | dH<sub>2</sub>O | 0.1 | 150-200 |
| HC-030031 | 50 | DMSO | 20 | 100 |
| A967079 | 10 | DMSO | 1 | 100 |
| RQ00203078 | 0.1** | dH<sub>2</sub>O | 1 | 100 |

## Acknowledgements

We thank Ardem Patapoutian and Ajay Dhaka for providing the Trpm8^EGFPf^ mouse line. The mouse Trpa1^-/-^ line was generated by Kevin Kwan and David Corey (Harvard Medical School). Ana Gómez del Campo performed the reanalysis of published single cell transcriptomic data. The authors are grateful to Laura Almaraz, Elvira de la Peña, Salvador Sala and Ana Gomis for comments. Mireille Tora and Victor Rodríguez Milan are acknowledged for excellent technical assistance. We thank Giovanna Exposito and Verona Villar for help with confocal imaging. During the course of this work, KGB was supported by the International PhD Fellowships Program “La Caixa”-Severo Ochoa, Call 2015”, PHO was supported by MICIU predoctoral fellowship BES-2017-080782. The work was funded by projects Generalitat Valenciana PROMETEO/2021/03, Spanish AEI projects PID2019-108194RB-I00 and PID2022140961OB-100/AEI/10.13039/501100011033, and co-financed by the European Regional Development Fund (ERDF) and the “Severo Ochoa” Program for Centers of Excellence in R&D (ref. CEX2021-001165-S).

## Authors’ contributions

KGB conducted calcium imaging, qPCR, histological studies and their analysis. PHO and EQ participated in the lung and stomach retrograde labelling experiments. FV conceived the study and was responsible for its coordination. Both, KGB and FV, prepared a draft of the manuscript. All authors read, corrected, and approved the final manuscript.

---

**Supplementary figure 1.**
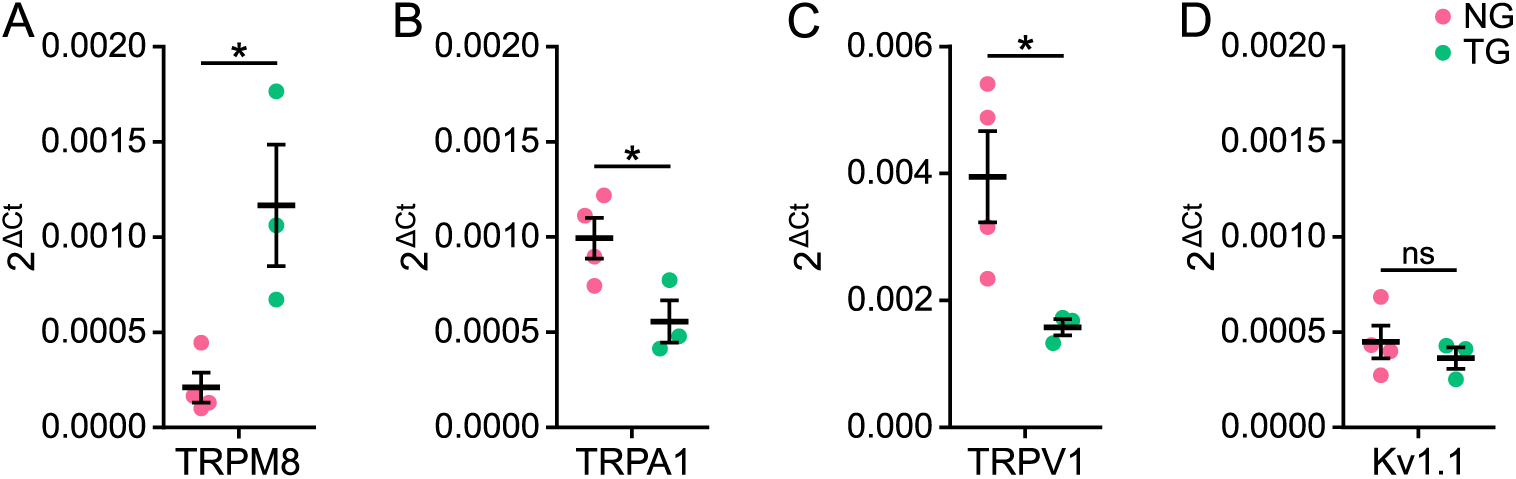
Differential TRP channel expression in VG and TG. Quantitative PCR data, normalized to two housekeeping genes (18S and β-actin), from RNA extracted from MACS purified neurons for TRPM8 (A), TRPA1 (B), TRPV1 (C) and Kv1.1 (D), (* p<0.05, unpaired t-test, n = 3-4 mice).

**Supplementary figure 2.**
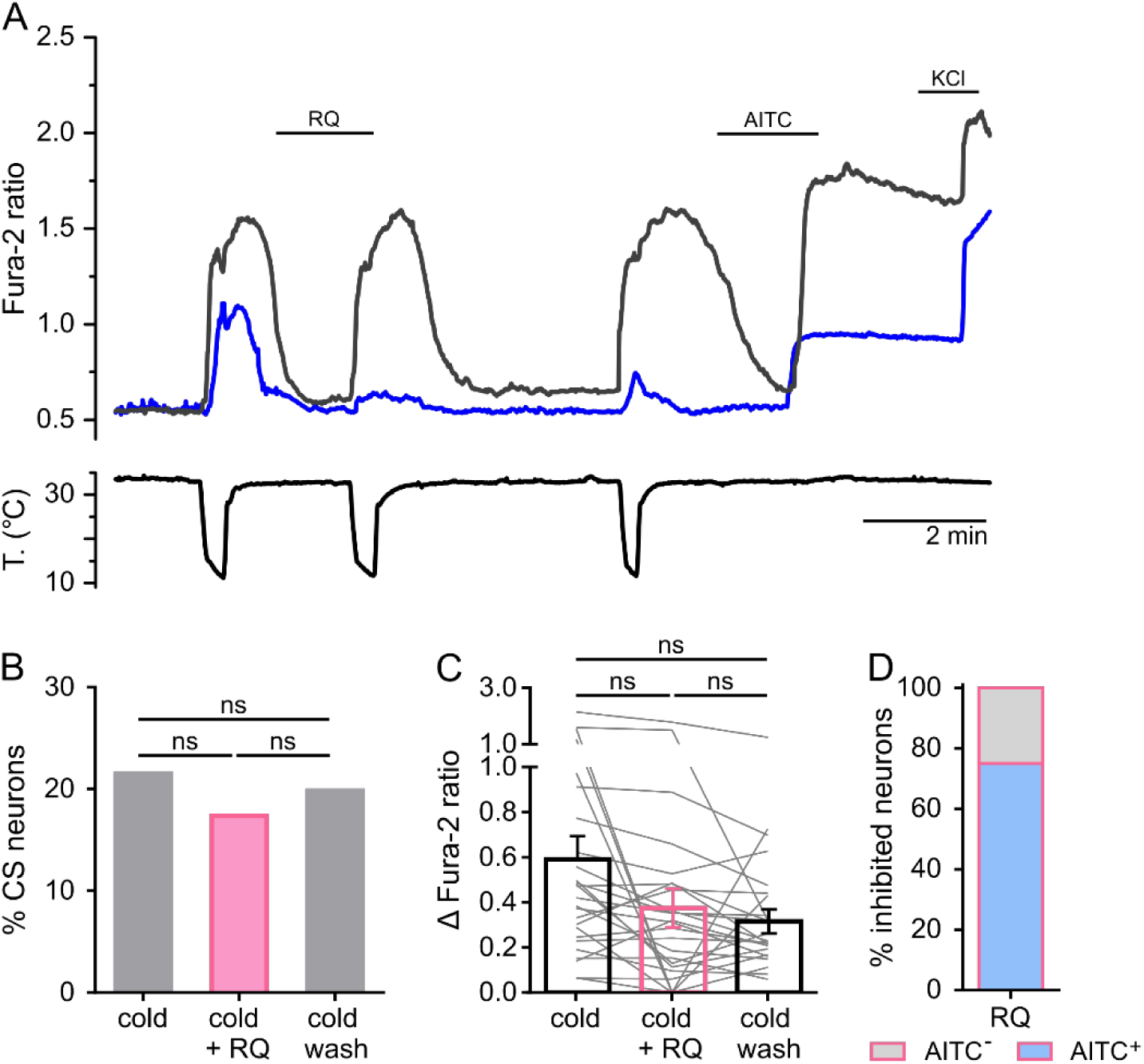
Pharmacological and genetic interrogation of cold sensitivity in VG neurons. (A) Exemplary traces of two vagal neurons from WT mice recorded simultaneously, showing differential effects of 1 µM RQ00203078 (RQ) on cold-evoked responses. Most neurons were insensitive to RQ (black trace) while a small fraction were inhibited (blue trace). Note that, regardless of sensitivity to RQ, both neurons were activated by 100 µM AITC and 50 mM KCl. (B) Percentage of cold responders to each of the three cold stimuli, Fisheŕs exact test, n= 121 cells. (C) Mean amplitudes to the three cold stimuli, light grey lines represent amplitudes of individual neurons, one-way ANOVA with Sidak’s post hoc, n= 26 cells. (D) Number of RQ-inhibited neurons that also responded to AITC (AITC^+^) or not (AITC^-^), n = 8 RQ-inhibited cells.

**Supplementary figure 3.**
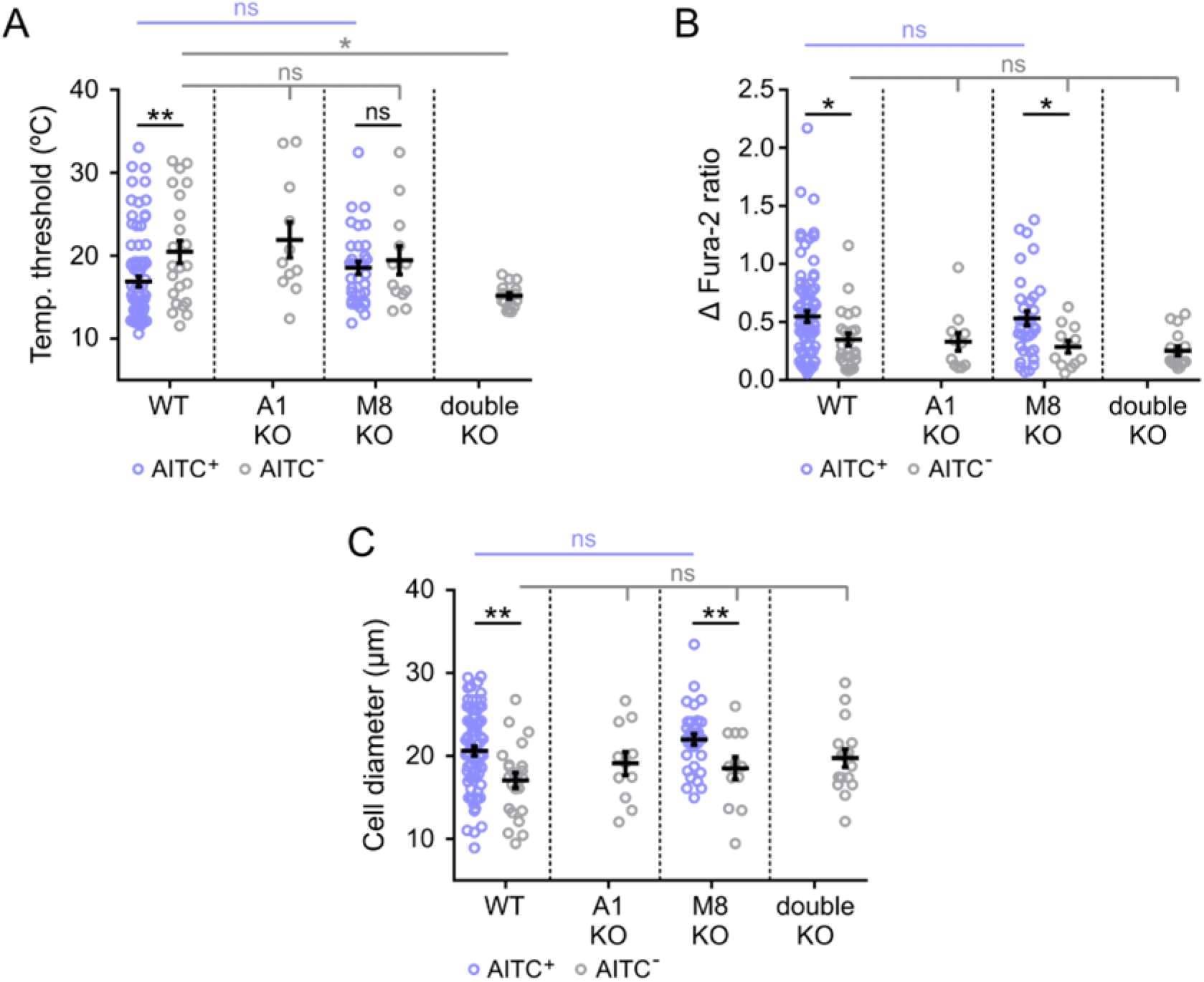
Differential functional and morphological properties of cold sensitive vagal neurons. Temperature thresholds (A), amplitude of cold-evoked responses (B), and cell diameters (C) for cold-sensitive VG neurons from different genotypes: blue circles represent AITC^+^ neurons (TRPA1-expressing), grey circles indicate AITC^-^ neurons. Note that AITC^+^ neurons have higher temperature thresholds, larger responses and larger diameters than AITC^-^ CS neurons. These neurons are absent in TRPA1 KO or double KO animals. One-way ANOVA with Tukeýs post hoc, * p<0.05, ** p<0.01, WT AITC+ n=79, WT AITC-n=23, A1 KO AITC+ n=0, A1 KO AITC-n=11, M8 KO AITC+ n=34, M8 KO AITC-n=12, dKO AITC+ n=0, dKO AITC-n=16 cells.

**Supplementary figure 4.**
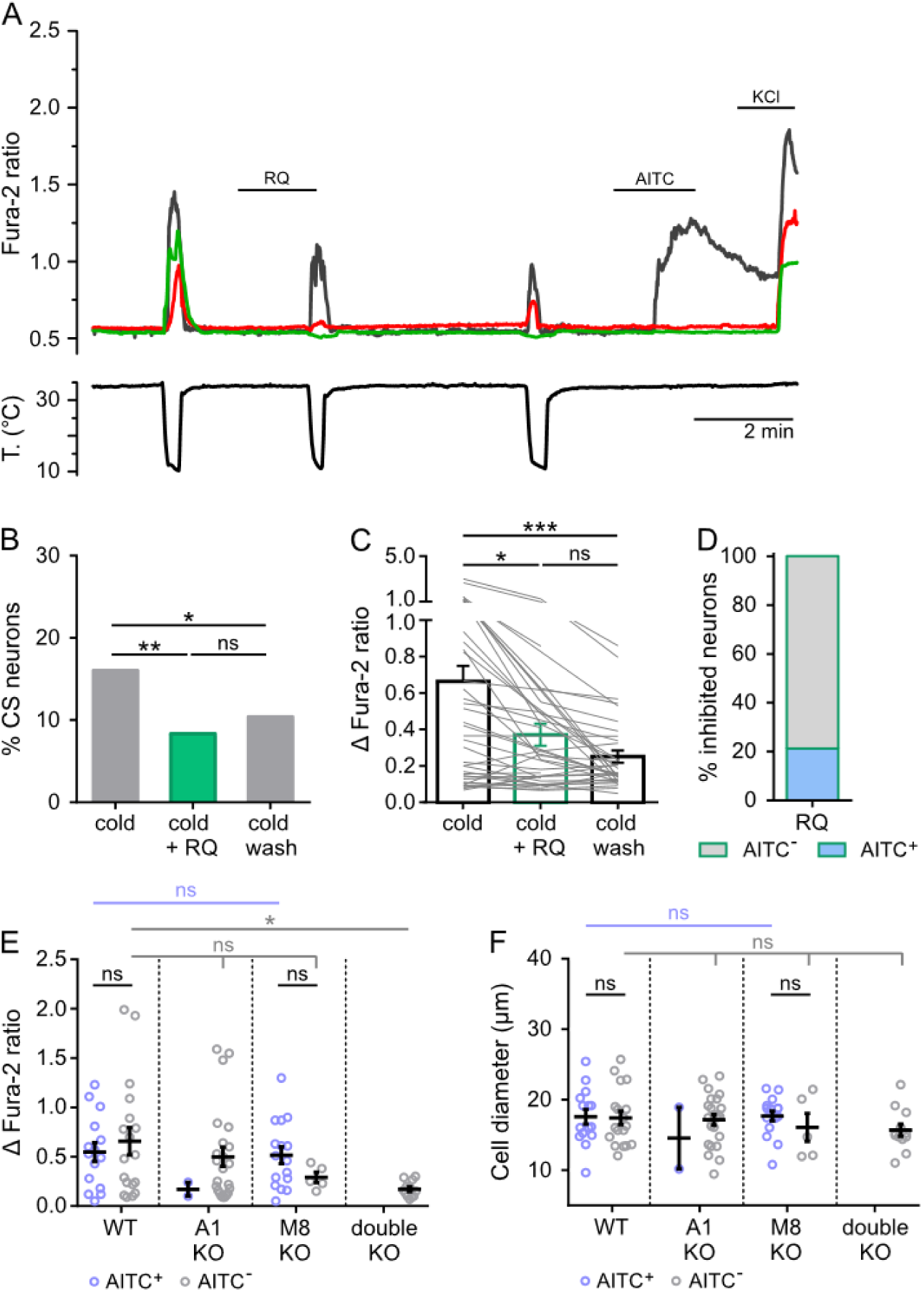
Pharmacological interrogation of CS neurons in TG. (A) Representative traces of neurons from WT mice that were exposed to a first cold ramp in control conditions, a second cold ramp in the presence of 1 µM RQ00203078 (RQ, 100 sec pre-incubation) and a third cold ramp 6-7 minutes after washing off RQ, 100 µM AITC, 50 mM KCl. (B) Number of cold responders to the three cold stimuli, Fisheŕs exact test, n= 337 cells. (C) Mean amplitudes to the three cold stimuli, light grey lines represent amplitudes of individual neurons, one-way ANOVA with Sidak’s post hoc, n= 54 cells. (D) Number of RQ-inhibited neurons that also responded to AITC (AITC^+^) or not (AITC^-^), n=33 RQ-inhibited cells. Amplitudes of cold responses (E) and cell diameter (F) for TG neurons from different genotypes, blue circles represent AITC^+^ neurons, grey circles indicate AITC^-^ neurons, one-way ANOVA with Tukeýs post hoc, * p<0.05, WT AITC^+^ n=15, WT AITC^-^ n=18, A1 KO AITC^+^ n=2, A1 KO AITC^-^ n=23, M8 KO AITC^+^ n=16, M8 KO AITC^-^ n=5, dKO AITC^+^ n=0, dKO AITC^-^ n=12 cells.

**Supplementary figure 5.**
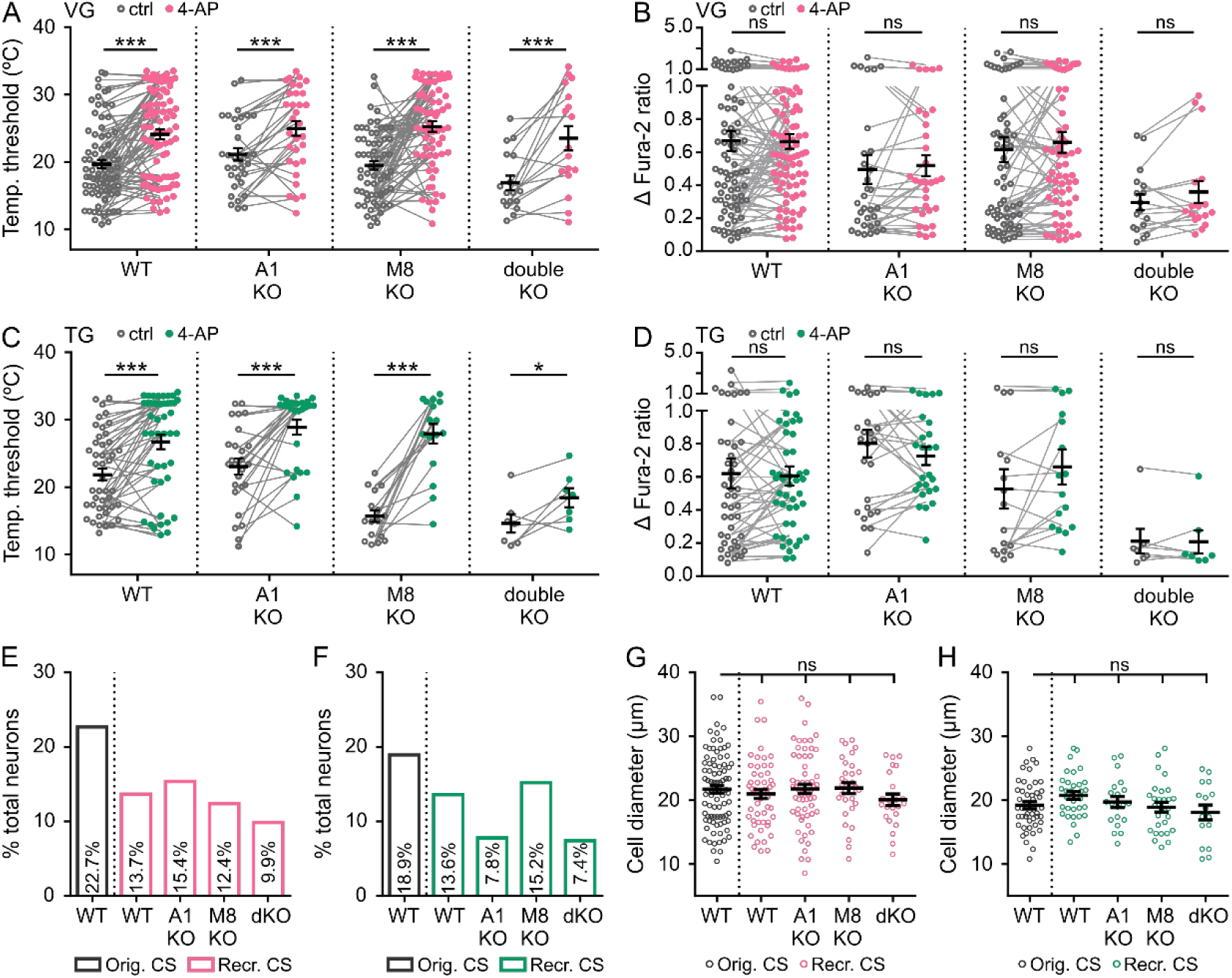
Characteristics of original and 4-AP-recruited CS neurons in VG and TG. (A-D) Temperature thresholds and amplitudes of original CS vagal (A,B) and trigeminal (C,D) neurons in control solution (open circles) or in the presence of 100 µM 4-AP (full circles), paired t-test, * p<0.05, *** p<0.001. (A,B) VG: WT n=82, A1 KO n = 32, M8 KO n = 63, dKO n = 17 cells. (C,D) TG: WT n=45, A1 KO n=26, M8 KO n=16, dKO n=7 cells. (E) Percentage of 4-AP recruited CS VG from different genotypes (pink bars) and of original CS neurons from WT mice (grey bar), WT n = 388, A1 KO n = 397, M8 KO n = 250, dKO n = 233 cells. (F) Percentage of 4-AP recruited CS TG from different genotypes (green bars) and of original CS neurons from WT mice (grey bar), original WT n = 243, A1 KO n = 257, M8 KO n = 184, dKO n = 216 cells. (G) Cell diameter of cold responses for recruited CS VG neurons from different genotypes (pink circles) and for original CS neurons from WT mice (grey circles), one-way ANOVA with Sidak’s post hoc, original WT n = 88, recruited WT n = 53, A1 KO n = 61, M8 KO n = 31, dKO n = 23 cells. (H) Cell diameter of cold responses for recruited CS TG neurons from different genotypes (green circles) and for original CS neurons from WT mice (grey circles), one-way ANOVA with Sidak’s post hoc, original WT n = 46, recruited WT n = 33, A1 KO n = 20, M8 KO n = 28, dKO n = 16 cells.

**Supplementary figure 6.**
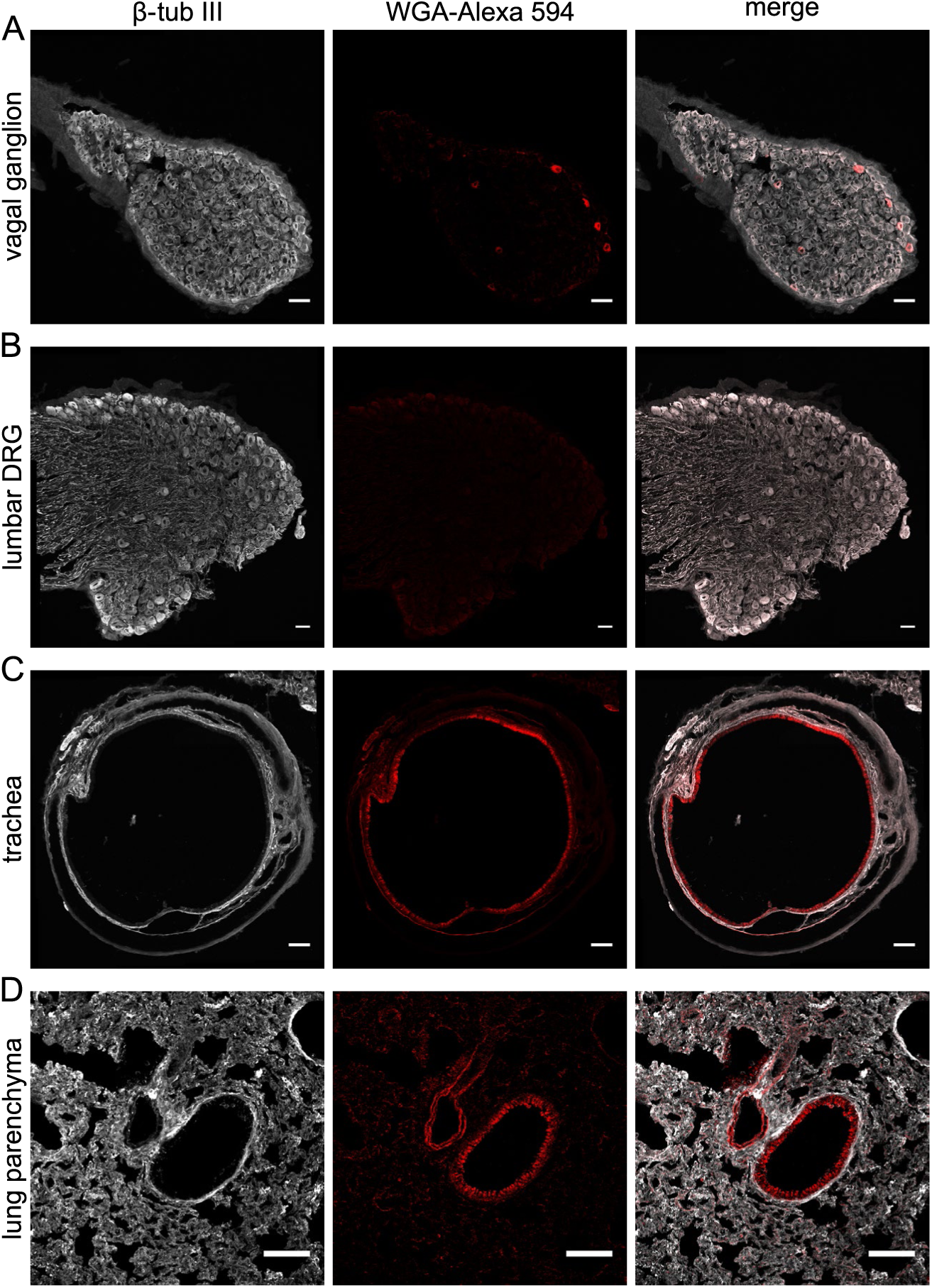
WGA-Alexa 594 staining can be found in VG, trachea, and lung parenchyma in fixed tissue sections. Confocal images of tubulin β III staining (left), WGA 594 fluorescence (middle) and the merged images (right) for VG (A), lumbar DRG (B), trachea (C) and lung parenchyma (D), scale bars 50 µm (A,B) and 100 µm, (C,D).

**Supplementary figure 7.**
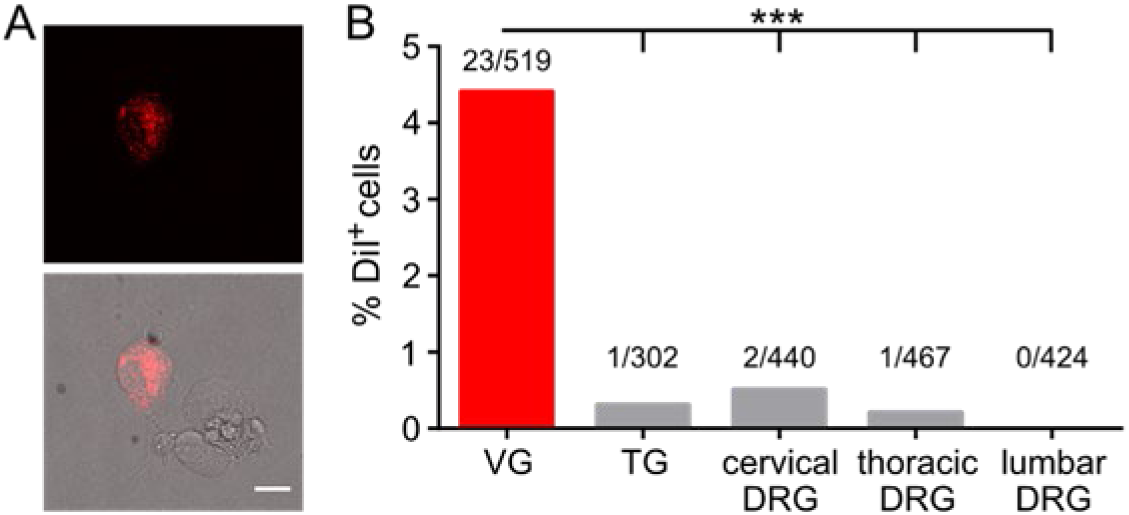
DiI retrograde labelling of airway-innervating neurons selectively labels vagal neurons. (A) Image of a fixed cultured DiI^+^ VG neuron under 550 nm (top) or transmitted light (bottom) exposure, scale bar 15 µm. (B) Percentage of DiI^+^ cells in different ganglia, number of DiI^+^/total neurons are stated above bars, cultures from two mice, *** p<0.001, Fisheŕs exact test compared to VG.

